# Embryonic gene transcription in the spiny mouse (*Acomys cahirinus*): an investigtion into the embryonic genome activation

**DOI:** 10.1101/280412

**Authors:** Jared Mamrot, David K. Gardner, Peter Temple-Smith, Hayley Dickinson

## Abstract

Our understanding of genetic mechanisms driving early embryonic development is primarily based on experiments conducted on mice, however translation of findings can be limited by physiological differences between mice and humans. To address this, we investigated whether the spiny mouse (*Acomys cahirinus*) is a closer model of early human embryonic development due to their more human-like endocrine profile. We therefore characterised the initiation of gene transcription in the spiny mouse embryo and compared the pattern of gene expression during the embryonic genome activation (EGA) with common mouse and human embryos. Naturally-mated spiny mouse embryos were obtained at the 2-cell, 4-cell and 8-cell stages of development (n=4 biological replicates per stage). RNA-Seq of these samples produced 709.1M paired-end reads in total. *De novo* assembly of reads was conducted using Trinity. Embryo-specific transcripts were extracted from the *de novo* assembly and added to the reference spiny mouse transcriptome. Transcription was first detected between the 2-cell and 4-cell stages for the majority of genes (n=3,428), with fewer genes first transcribed between the 4-cell and 8-cell stages (n=1,150). The pattern of gene expression in spiny mouse embryos during this period of development is more human-like than common mouse embryos. This is the first evidence the spiny mouse may provide a more suitable model of human embryonic development. The improved reference *Acomys cahirinus* transcriptome is publically accessible, further increasing the value of this tool for ongoing research. Further investigation into early development in the spiny mouse is warranted.

---

The spiny mouse (*Acomys cahirinus*) is a small rodent native to regions of the Middle East and Africa (Nowak, 1999; Wilson and Reeder, 2005). It displays several unique physiological traits, including the capacity to regenerate skin without fibrotic scarring (Gawriluk et al., 2016; Seifert et al., 2012; Simkin et al., 2017) and a human-like endocrine profile, including cortisol as the primary glucocorticoid and production of DHEA (Quinn et al., 2013; Quinn et al., 2016). Unlike most eutherian mammals, the Egyptian spiny mouse (‘common spiny mouse’) has a menstrual cycle (Bellofiore et al., 2017). It is the only known species of rodent that menstruates and there are important differences in early embryonic development and implantation in menstruating species compared to those with an oestrus cycle such as mice, rats, cows, sheep and pigs (Brevini et al., 2006; Brosens et al., 2009; Emera et al., 2012; Graf et al., 2014; Memili & First, 2000; Niakan et al., 2012; Telford et al., 1990). Differences such as the polarity of apical attachment and cellular communication between the embryo and the endometrium can be identified before an embryo has implanted, and we may better understand the underlying mechanisms determining pregnancy success or failure by using a menstruating mammal to model human embryonic development in place of the mouse (*Mus musculus*) (Aplin & Ruane 2017; Brosens et al., 2009; Wang & Dey, 2006; Whitby et al., 2017). Little is known about early development in the spiny mouse, however preliminary evidence suggests it may overcome limitations of other species for modelling embryo development in humans.

One publication exists on spiny mouse embryo development in which the authors established methods for producing and culturing spiny mouse embryos *in vitro* (Pasco et al., 2012). One of the key challenges identified in this study was the presence of a ‘4-cell block’, with embryos unable to develop past 4-cells when cultured outside of the reproductive tract. Embryos developed *in vivo* obtained at the 8-cell stage are able to be cultured successfully *in vitro* through to the implantation stage, however the timing of the cell block is an example of differences between *Mus musculus* and spiny mouse embryos at the molecular level (Taft, 2008). Mouse embryos exhibit a 2-cell block when exposed to inadequate culture conditions, whereas human embryos exhibit a 4- to 8-cell block (Braude et al., 1988; Goddard & Pratt, 1983). The cellular environment is a major influence on gene expression in preimplantation embryos (Gardner & Kelley, 2017; Mantikou et al., 2017); characterising gene expression profiles during embryogenesis may therefore help direct future research efforts to overcome the 4-cell block in the spiny mouse and promote its use as a model of human embryo development.

Embryogenesis is a complex process regulated by diverse, interdependent physiological mechanisms. Successful development from a single cell (zygote) to live offspring requires coordinated changes in cell cycle, chromatin state, DNA methylation and genome conformation. Cellular machinery for transcription and translation must be successfully assembled, and transcription of the incipient genome must take place. Failure to successfully attain any of these developmental milestones results in death of the organism. The first major developmental transition in eukaryotic embryos is the maternal-to-zygotic transition (MZT), which involves clearance of maternally-inherited transcripts and transcription of the newly formed embryonic genome (the embryonic genome activation, ‘EGA’) (Ivanova et al., 2017; Schier, 2007; Tadros & Lipshitz, 2009). Next Generation Sequencing (NGS) can be used to comprehensively characterise this event. In mammals the MZT typically occurs between the 1-cell stage and 16-cell stages of development, however the timing and pattern of embryonic gene expression is species-specific (Tadros & Lipshitz, 2009). In mice, the MZT occurs predominantly between the 1-cell and 4-cell stages, with the EGA beginning at the 2-cell stage (Flach et al., 1982; Wang and Dey, 2006). In comparison, in human embryos the EGA begins at the 4- to 8-cell stage (Braude et al., 1988; Tesarik et al., 1988). The timing of these events coincides with the timing of the ‘cell block’ previously described in these species (Braude et al., 1988; Goddard & Pratt, 1983).

Recent studies on human embryos have identified ∼150 genes upregulated from the oocyte to 1-cell stage, followed by ∼1,000 genes upregulated from the 2-cell to 4-cell stage (Xue et al., 2013; Yan et al., 2013), and >2,500 genes first transcribed between the 4-cell and 8-cell stage. The specific genes activated during each stage of the EGA have been shown to differ significantly between mice and humans, with reports of only ∼40% concurrence between these two species (Heyn et al., 2014; Xie et al., 2010). Despite this, expression of specific genes driving the EGA are similar between humans and mice, and the overall pattern of transcription follows a similar pattern in mammals such as the cow, sheep, rabbit and other primates, occurring in ‘waves’ with different genes transcribed at different timepoints (Dobson et al., 2004; Taylor et al., 1997; Tesari’k et al., 1987; Vassena et al., 2011). Although the pattern of EGA in rodents is similar to humans, conspicuous differences exist (Christians et al., 1994; Crosby et al., 1988; De Sousa et al., 1998; Frei et al., 1989; Schramm and Bavister, 1999; Telford et al., 1990) and the search for a more suitable model continues.

The aims of this study were to characterise early embryonic gene expression, identify the period of EGA, and to compare the pattern of global gene expression in the spiny mouse to existing datasets from human and mouse embryos. We hypothesise the EGA in the spiny mouse embryo will more closely reflect the EGA in human embryos than mouse embryos.

## METHODS

### Sample preparation and RNA sequencing

Embryos were collected from female spiny mice (n=12) in accordance with the Australian Code of Practice for the Care and Use of Animals for Scientific Purposes with approval from the Monash Medical Centre Animal Ethics Committee. Female dams were staged from delivery of their previous litter (spiny mice conceive their next litter approximately 12h postpartum) and culled at specific time-points for embryo retrieval at the required stage: 2-cell at 48h postpartum (n=4), 4-cell at 52h postpartum (‘early’ 4-cell; n=2) or at 68h postpartum (‘late 4-cell’; n=2), and 8-cell at 72h postpartum (n=4). Embryos were flushed from the excised reproductive tract using warmed G-MOPS PLUS handling medium containing 5 mg/ml human serum albumin (Vitrolife, Göteborg, Sweden), washed through warmed sterile Ca^2+^/Mg^2+^-free PBS three times using sterile pulled glass pipettes, and grouped into biological replicates (n=4 for each stage: 12 samples total). Embryos were snap frozen using liquid nitrogen in a minimal volume of cell lysis solution (∼1μl) comprised of lysis buffer, dithiothreitol (DTT) and RNase inhibitors per NuGEN SoLo RNA-Seq kit (NuGEN Technologies, Inc; San Carlos, CA, USA). To reduce the impact of embryo collection and freezing on gene transcription this process was conducted as quickly as possible: embryos were snap-frozen in lysis solution using liquid nitrogen and stored at −80°C in less than 5 minutes post-mortem.

To aid lysis, two freeze-thaw cycles were conducted on a slurry of dry ice and ethanol prior to library preparation. Samples were then processed per the Nugen SoLo protocol (version M01406v3; available from NuGEN). After ligation of cDNA, qPCR was performed on all samples to determine the number of amplification cycles required to ensure that amplification was in the linear range. Based on these results, each sample was amplified using 24 cycles. Final libraries were quantitated by Qubit and size profile determined by the Agilent Bioanalyzer.

Custom ‘AnyDeplete’ rRNA depletion probes were designed and produced by NuGEN Technologies, Inc (San Carlos, CA, USA) using rRNA sequences from the spiny mouse transcriptome (Mamrot et al., 2017). Prior to use, efficacy and off-target effects of the rRNA depletion probes were examined *in silico* by NuGEN. Samples were loaded using c-Bot (200pM per library pool) and run on 2 lanes of an Illumina HiSeq 3000 8-lane flow-cell. PhiX spike-in was not used directly due to incompatibility with the custom rRNA depletion probes, however it was incorporated into other lanes of the same run. RNA-Seq data (100bp, paired-end reads) were uploaded to the NCBI under Bioproject PRJNA436818 (SRA : SRP133894).

The quality of RNA-Seq reads was assessed using FastQC v0.11.6 (https://github.com/s-andrews/FastQC; 50f0c26), with MultiQC v1.4 (https://github.com/ewels/MultiQC; baefc2e) report available from Github (https://github.com/jpmam1) (Ewels et al., 2016). Adapter sequences were trimmed from the reads using trim-galore v0.4.2 (https://github.com/FelixKrueger/TrimGalore; d6b586e), implementing cutadapt v1.12 (https://github.com/marcelm/cutadapt; 98f0e2f). Reads with a quality scores lower than 20 and read pairs in which either forward or reverse reads were trimmed to fewer than 35 nucleotides were discarded. Further trimming was conducted using Trimmomatic v0.36 (http://www.usadellab.org/cms/index.php?page=trimmomatic) with settings “`LEADING:3 TRAILING:3 SLIDINGWINDOW:4:20 AVGQUAL:25 MINLEN:35`” (Bolger et al., 2014). Nucleotides with quality scores lower than 3 were trimmed from the 3’ and 5’ read ends. Reads with an average quality score lower than 25 or with a length of fewer than 35 nucleotides after trimming were removed. Error correction of trimmed reads was performed using Rcorrector v1.0.2 (https://github.com/mourisl/Rcorrector; 144602f) (Song & Florea, 2015). FastQC was used to assess the improvement in read quality after trimming adapter removal; MultiQC report is available from Github (https://github.com/jpmam1).

### *De novo* transcriptome assembly and read alignment

Error corrected reads were assembled using Trinity v2.4.0 (https://github.com/trinityrnaseq/trinityrnaseq; 1603d80) with settings “`--max_memory 400G, -- CPU 32 and --full_cleanup`” (Haas et al., 2013). Assembly statistics were computed using the TrinityStats.pl script from the Trinity package and are provided in Table S1. All reads were aligned to the assembled ‘embryo’ transcriptome using Bowtie2 v2.2.5 (https://github.com/BenLangmead/bowtie2; e718c6f) with settings: “`--end-to-end, --score-min L,-0.1,-0.1, --no-mixed, --no-discordant, -k 100, -X 1000, --time, -p 24`” (Langmead & Salzberg, 2012).

Read-supported contigs were identified within the ‘embryo’ transcriptome assembly using samtools “idxstats” v1.5. (https://github.com/samtools/samtool; f510fb1) (Li et al., 2009). Read support was defined as >=1 reads aligned. Read-supported contigs from the embryo-specific assembly were added to the reference spiny mouse transcriptome assembly previously described by Mamrot et al. (2017) and samples were aligned to this ‘updated’ transcriptome using Bowtie2 with settings “`--end-to-end, -- score-min L,-0.1,-0.1, --no-mixed, -- no-discordant, -k 100, -X 1000, --time, -p 24`”.

Trinity contigs were aligned to the UniProtKB/SwissProt protein sequence database (ftp://ftp.uniprot.org/pub/databases/uniprot/current_release/knowledgebase/complete/uniprot_sprot.fasta.gz accessed 14th October 2017) using BLASTx v2.5.0+ (ftp://ftp.ncbi.nlm.nih.gov/blast/executables/blast+/2.5.0/) (Altschul et al., 1997). Confident BLAST hits were retained, transcripts were annotated using the single-best hit based on e-value, and Gene Ontology (GO) terms were obtained for further analysis. Trinity-normalized reads were aligned to the NCBI nr protein database (ftp://ftp.ncbi.nih.gov/blast/db/FASTA/nr.gz accessed 10th February 2018) using DIAMOND v0.9.17 blastx (Buchfink et al., 2015) with taxonomic and functional annotation of reads aligning to eukaryotic and prokaryotic lineages conducted using MEGAN6 Community Edition v6.10.10 (https://ab.inf.uni-tuebingen.de/software/megan6) (Huson et al., 2016). MEGAN6 files were accessed using MeganServer v1.0.1 (https://ab.inf.uni-tuebingen.de/software/meganserver) (Beier et al., 2017). Reads were also aligned to mouse and human RefSeq rRNA sequences (accessions: NR_003279.1, NR_003278.3, NR_003280.2, NR_046144.1, NR_003285.2, NR_003287.2, NR_003286.2, X71802.1) and mouse tRNAs within the GtRNAdb database (http://gtrnadb.ucsc.edu/genomes/eukaryota/Mmusc10/mm10-tRNAs.fa) (Chan & Lowe, 2016).

### Transcript clustering and differential gene expression

Read alignments to the updated transcriptome generated using Bowtie2 were clustered with Corset v1.0.7 (https://github.com/Oshlack/Corset; cf4d4fb) to reduce the impact of redundant transcripts and transcript isoforms when assessing gene expression (Davidson and Oshlack, 2015). Variance in RNA-Seq data was explored using Varistran v1.0.3 (https://github.com/MonashBioinformaticsPlatform/varistran; ff90258), which implements Anscombe’s variance stabilizing transformation (1948) to equalize noise across all samples before assessing gene expression levels (Harrison, 2017). Differential gene expression was explored using the Degust web application (http://degust.erc.monash.edu/). Further investigation was conducted using EdgeR v3.20.8 (https://bioconductor.org/packages/release/bioc/html/edgeR.html) (Robinson et al., 2010). Correlations were calculated using Corrplot v0.83 (https://github.com/taiyun/corrplot; d7ba847) (Wei & Simko, 2017). Confidence bounds for effect sizes were calculated using TopConfects v1.0.1 (https://github.com/pfh/topconfects; 43cd006) (Harrison, 2018).

### Profiling gene expression during the EGA

Gene expression data were accessed for mouse and human embryos from the NCBI Gene Expression Omnibus (GEO) project GSE44183 (accessed 22/02/2018) (Xue et al., 2013). This dataset contains both human and mouse embryos collected at the same developmental stages (mouse: 3X2-cell, 3X4-cell, 3X8-cell; human: 3X2-cell, 4X4-cell and 10X8-cell). Gene expression profiles were generated from Log2 fold changes in Fragments Per Kilobase of transcript per Million mapped reads (FPKM) extracted from expression matrices provided by the authors (ftp://ftp.ncbi.nlm.nih.gov/geo/series/GSE44nnn/GSE44183/suppl/). All figures were produced using R software v3.4.0 and GraphPad Prism 7.

## RESULTS

### RNA sequencing and quality control

In total, 701.9 million reads passed filtering across 12 samples (Table 1) with a relatively high proportion of >Q30 reads (95.2%). The error rate was 0.2% (expected <0.5%) and phasing/prephasing was 0.13/0.08 (expected <0.4/<0.2), indicating high-quality sequencing with minimal technical errors. Read error correction resulted in 266 million repairs (∼0.1% of all nucleotides). Quality metrics obtained using FastQC before and after read processing are available from Github: https://rawgit.com/jpmam1/multiQC_reports/master/pre-trimming_multiqc_report.html https://rawgit.com/jpmam1/multiQC_reports/master/post-trimming_multiqc_report.html.

**Table 1:** Summary of paired-end (100bp) Illumina sequencing output for each sample

| ULN | Sample name | Biosample | I7 Index | Mean Library Size (bp)* | Identifier sequence | Reads Passed Filter (Million) |
| --- | --- | --- | --- | --- | --- | --- |
| 17---04796 | Sample1 ("2cell_A") | SRR6804613 | C02 | 338 | GACTACGA | 58.1 |
| 17---04797 | Sample2 ("2cell_B") | SRR6804612 | D02 | 318 | ACTCCTAC | 62.1 |
| 17---04798 | Sample3 ("2cell_C") | SRR6804607 | E02 | 335 | CTTCCTTC | 61.9 |
| 17---04799 | Sample4 ("2cell_D") | SRR6804606 | F02 | 338 | ACCATCCT | 59.2 |
| 17---04800 | Sample5 ("4cell_A") | SRR6804609 | G02 | 338 | CGTCCATT | 59.6 |
| 17---04801 | Sample6 ("4cell_B") | SRR6804608 | H02 | 354 | AACTTGCC | 57.2 |
| 17---04802 | Sample7 ("4cell_C") | SRR6804611 | A03 | 342 | GTACACCT | 53.2 |
| 17---04803 | Sample8 ("4cell_D") | SRR6804610 | B03 | 343 | ACGAGAAC | 54.5 |
| 17---04804 | Sample9 ("8cell_A") | SRR6804617 | C03 | 323 | CGACCTAA | 64.4 |
| 17---04805 | Sample10 ("8cell_B") | SRR6804616 | D03 | 321 | TACATCGG | 55.9 |
| 17---04806 | Sample11 ("8cell_C") | SRR6804615 | E03 | 327 | ATCGTCTC | 56.8 |
| 17---04807 | Sample12 ("8cell_D") | SRR6804614 | F03 | 365 | CCAACACT | 59.8 |
| ----- |  |  |  |  |  |  |
| Total reads |  |  |  |  |  | 701.9 |

\* The expected sample library size range was ~320-360 bp.

### *De novo* transcriptome assembly and read alignment

All trimmed and error-corrected reads were assembled into an ‘embryo’ transcriptome (assembly metrics: Table S1) to detect transcripts specific to early development not present in our reference spiny mouse transcriptome. The proportion of reads mapping to this embryo-specific transcriptome, the number of unique reads per sample, and proportion of reads from each sample aligned to human/mouse rRNA sequences are shown in Figure 1 (no reads aligned to the tRNA database). Transcripts from the embryo assembly were aligned to the UniProtKB / SwissProt protein database using BLASTx: ∼70% of transcripts aligned to *Mus musculus, Homo sapiens* and *Rattus norvegicus*, and ∼30% aligned to other eukaryotic and prokaryotic taxa (interactive summary: https://public.flourish.studio/visualisation/20088/).

**Figure 1:**
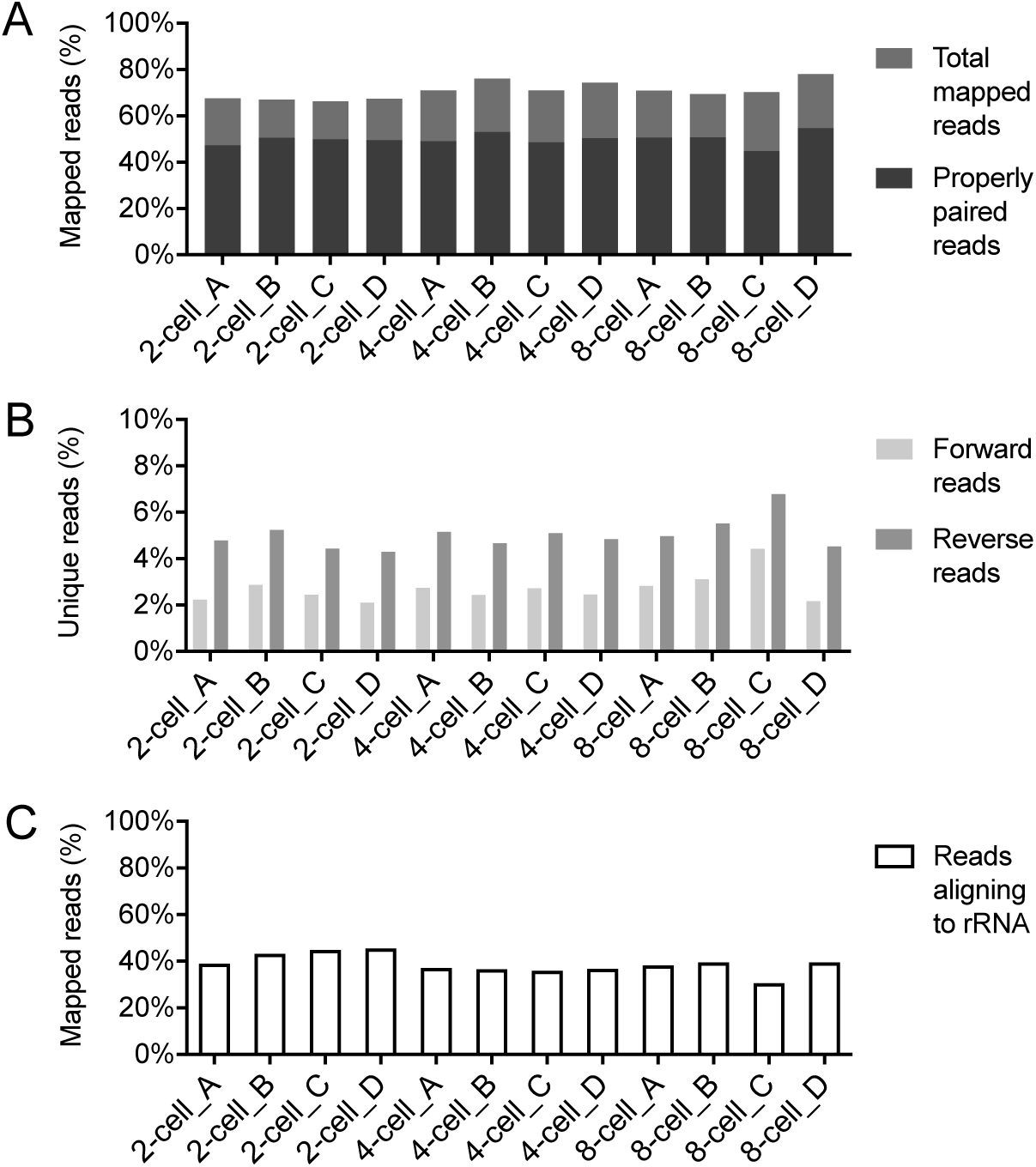
(A) Proportion of reads mapping to the ‘embryo’ spiny mouse transcriptome assembly. “Properly paired reads” both align to the same transcript, “Total mapped reads” represent either forward or reverse reads mapped to a transcript. (B) The proportion of unique reads per sample, and (C) reads mapping to human / mouse RefSeq rRNA sequences.

### Clustering and differential gene expression

Transcripts from the *de novo* assembly (n=595,435) were clustered together based on read mapping to form 309,543 representative gene clusters (from here on referred to as ‘genes’). This clustering facilitated use of gene-level methods for quantification and analysis. Average read count for each sample library and hierarchical clustering of samples based on average gene abundance in counts-per-million (cpm) are shown in Figure 2A (further sample correlations are shown in Figure S3). Hierarchical clustering revealed clear differentiation between 2-cell embryos and the 4-cell/8-cell embryos, with less clear differentiation between 4-cell and 8-cell embryos (Figure 2B).

**Figure 2:**
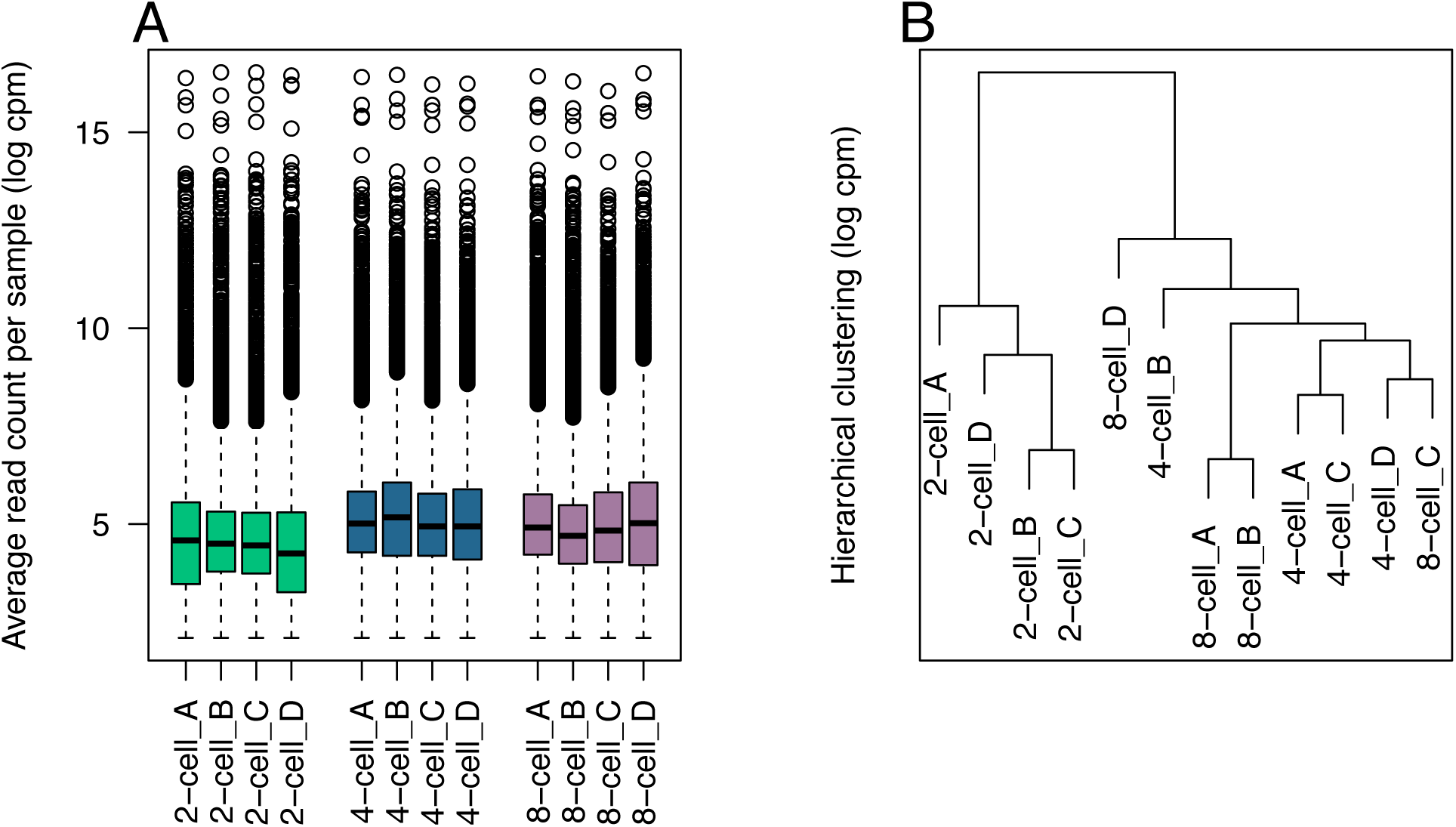
(A) Average read count per sample and (B) hierarchical clustering of samples based on gene abundance in each library (cpm = count per million).

Application of Anscombe’s variance stabilizing transformation tempered dispersion across all samples (average dispersion = 0.0784). Library sizes before and after ‘trimmed mean of M’ (TMM) normalization (Robinson & Oshlack, 2010) are listed in Supplementary Table 2. Read counts were clustered in two dimensions to examine group differences in gene abundance. Multi-dimensional scaling (MDS) analysis suggests two of the samples (“2cell_C” and “8cell_A”) have atypical profiles compared to the other samples (Figure 3).

**Figure 3:**
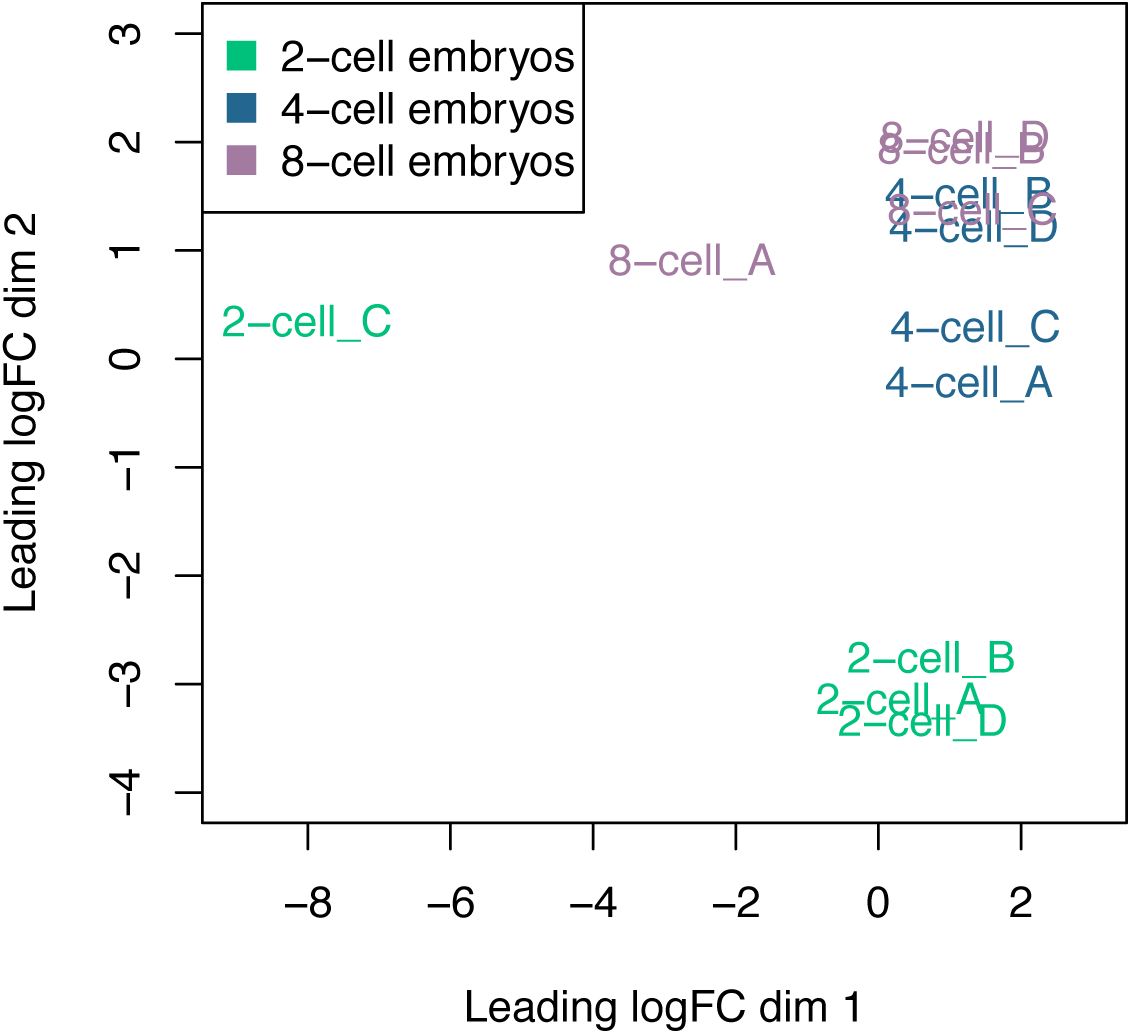
MDS plot illustrating differences in average gene expression between samples for the top 500 differentially-expressed genes (n=12).

Further investigation revealed significant differences between samples ‘2-cell_C’ and ‘8-cell_A’ compared to the other 10 samples (Figure 4). BLASTx alignment of differentially expressed (DE) genes to the UniProtKB/SwissProt database revealed evidence of contamination, with >800 bacteria-associated genes highly expressed in these two samples and no expression detected in the other samples (Table S3). Metatranscriptomic analysis of read alignments to the NCBI nr database confirmed significant prokaryotic contamination in samples ‘2-cell_C’ and ‘8-cell_A’ (Figure S1). These samples were not able to be salvaged due to the level of contamination and were excluded from further analysis, reducing statistical power (Figure S2). With these two samples removed the gene expression profiles of spiny mouse embryos are comparable to other mammals at this stage of development (Figure 5).

**Figure 4:**
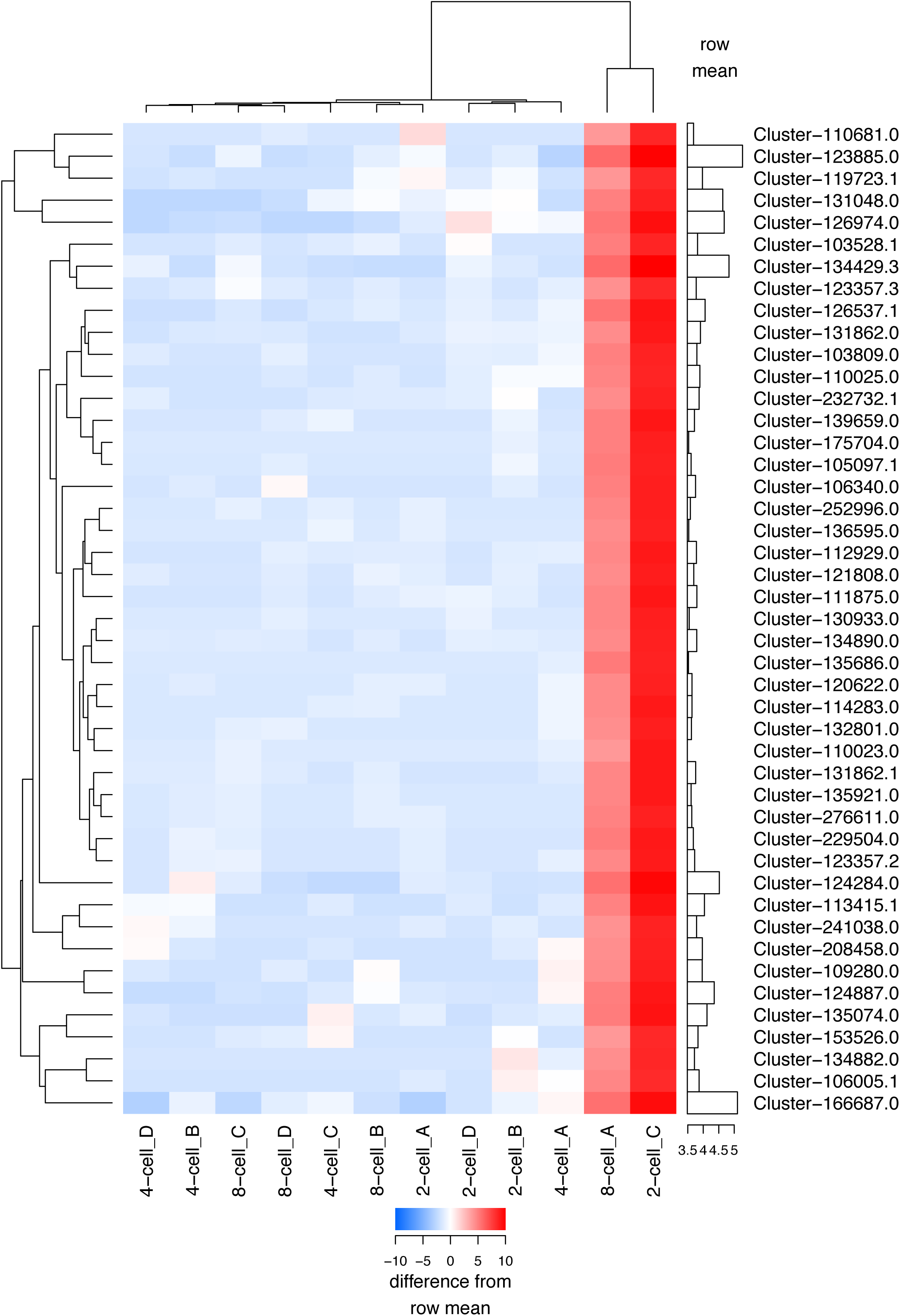
Heatmap of top 50 differentially expressed genes (n=12). Gene expression in samples 2-cell_C and 8-cell_A is highly abnormal due to the presence of prokaryotic contamination.

**Figure 5:**
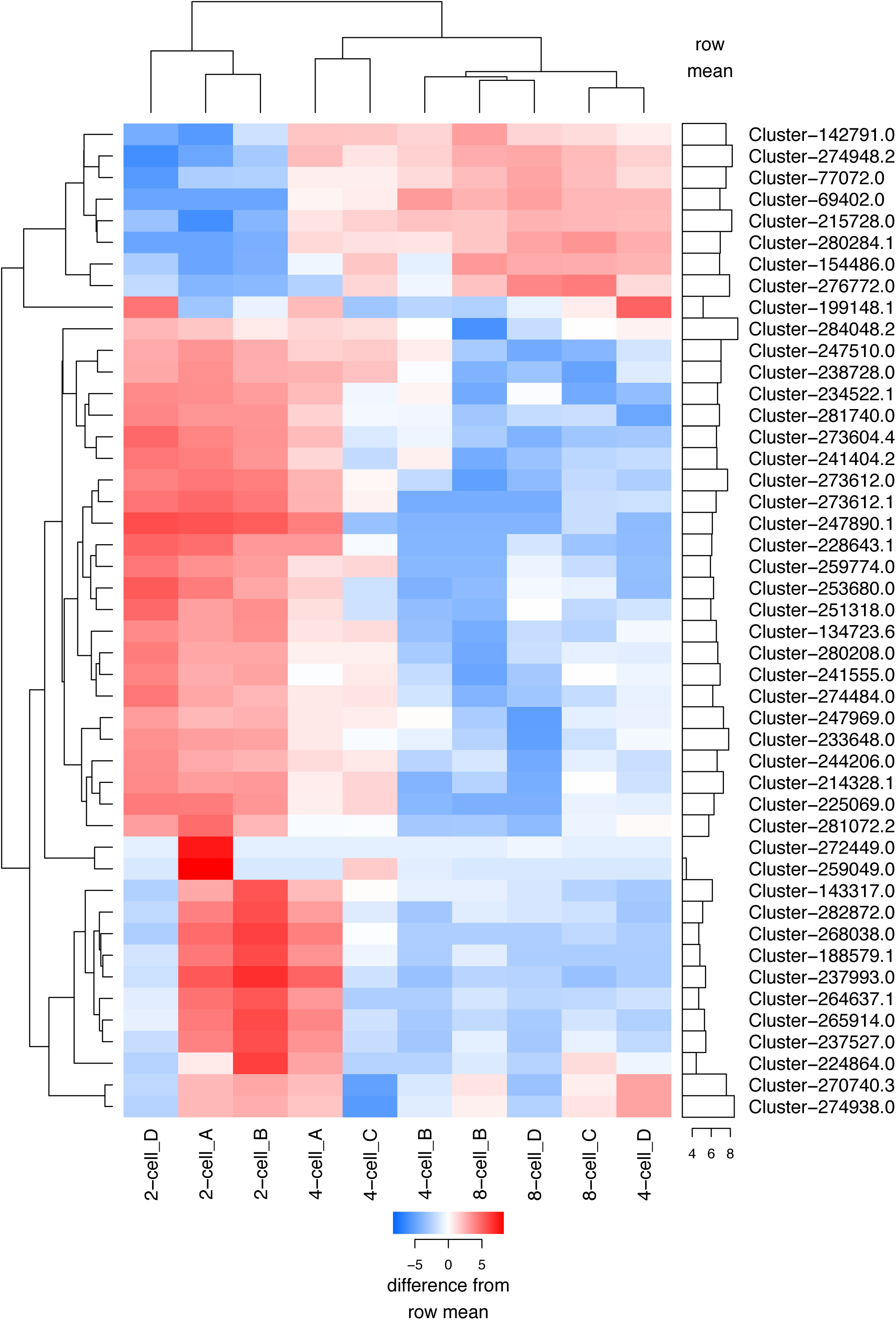
Heatmap of the top 50 differentially expressed genes with samples 2-cell_C and 8-cell_A excluded from analysis (n=10).

Alignments from the 10 uncontaminated samples to the ‘embryo’ transcriptome were re-examined. All transcripts with >=1 reads aligned were extracted (54,660 read-supported contigs in total; 441.23 Mb of sequence data) and added to the ‘reference’ transcriptome assembled by Mamrot et al. (2017). Alignment, clustering and gene expression analysis were performed against the ‘updated’ reference transcriptome. Transcripts from the updated assembly (n=2,274,638) were clustered based on read mapping using Corset; the number of gene clusters produced using the updated reference assembly (n=253,449) was fewer than the number produced using the *de novo* ‘embryo’ assembly (n=309,543). Exclusion of contaminated samples resulted in stronger correlations within developmental stages (Figure 5, Figures S4 & S5) and increased delineation between developmental stages, with ‘early’ 4-cell embryo samples (4-cell_A and 4_cell_C) clustering more closely to the 2-cell embryos, and the ‘late’ 4-cell samples (4-cell_B and 4_cell_D) clustering more closely to the 8-cell embryos (Figures 5 and 6).

**Figure 6:**
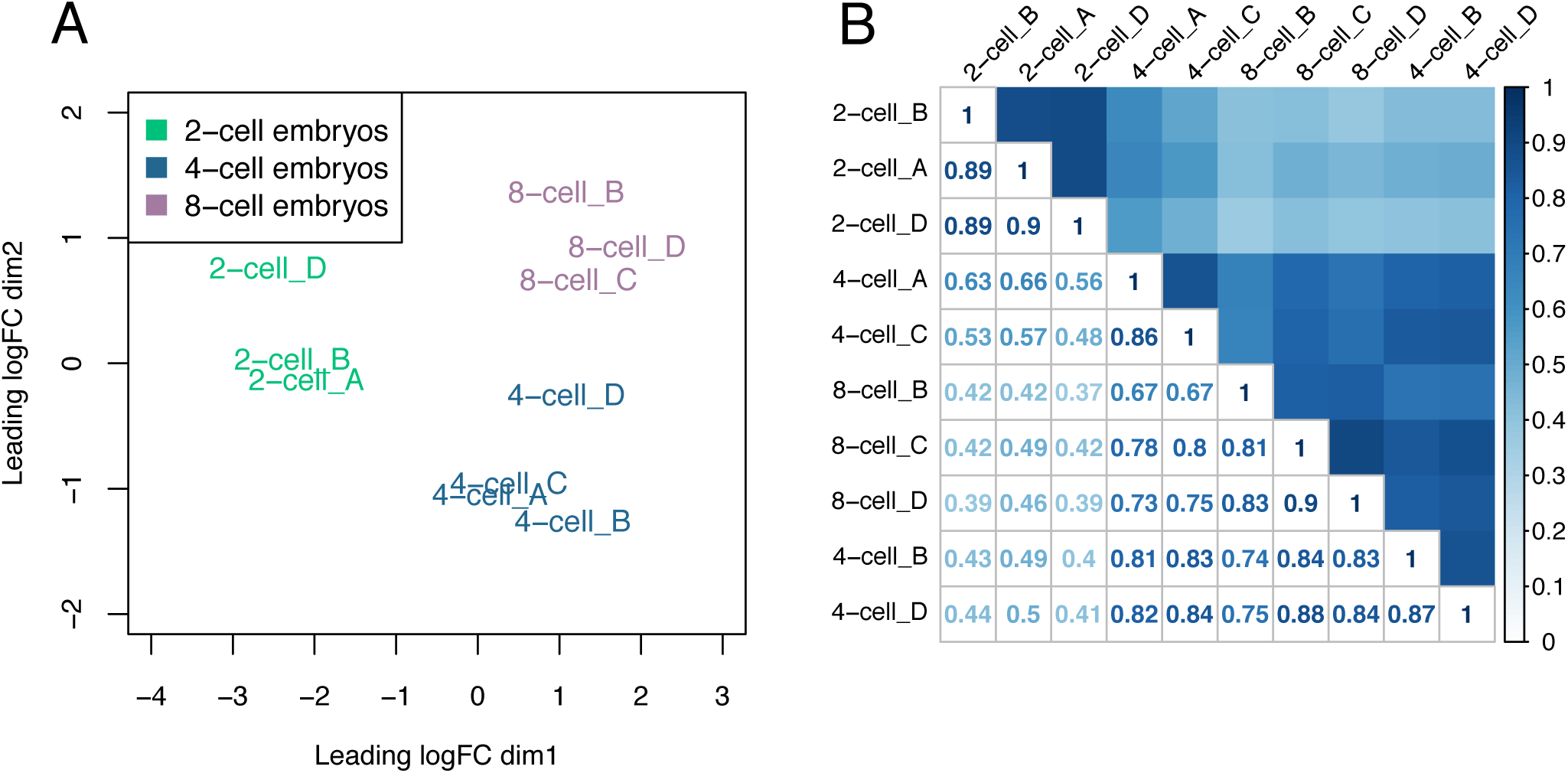
(A) MDS plot for top 1000 genes in remaining uncontaminated samples after reanalysis (n=10), and (B) corresponding Spearman rank correlations of gene abundance (n=10). Samples 4-cell_A and 4-cell_C were collected at 52h post-partum (“4-cell early”), and samples 4-cell_B and 4-cell_D were collected at 68h postpartum (“4-cell late”).

Fit of the negative binomial distribution to gene counts (Figures S6 & S7), biological coefficient of variation / quasi-likelihood dispersion (Figure S8), and mean-difference of each sample against combined samples (Figure S9) support the use of quasi-likelihood F-tests to determine differential expression. In total, differentially expression was detected in 3,428 genes between the 2-cell and 4-cell stages and 1,150 genes between the 4-cell and 8-cell stages of embryo development in the spiny mouse (Figures 7 and 8).

**Figure 7:**
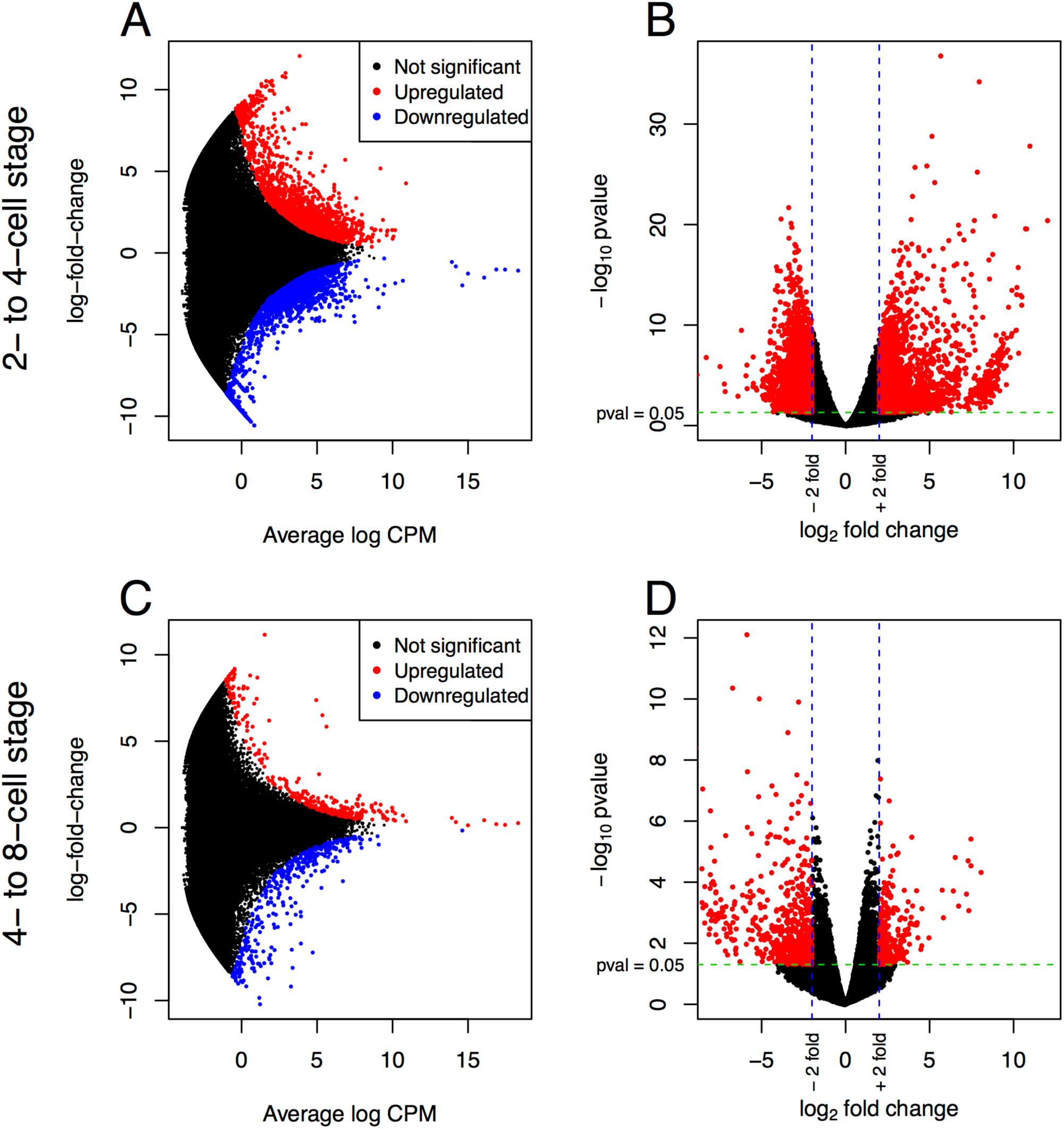
Differentially expressed genes between the 2-cell and 4-cell stages of development (A & B) and the 4-cell and 8-cell stages of development (C & D). Coloured dots represent individual differentially expressed genes. Smear plots: FDR<0.05 (A & C). Volcano plots: p-value <0.05 (B & D).

**Figure 8:**
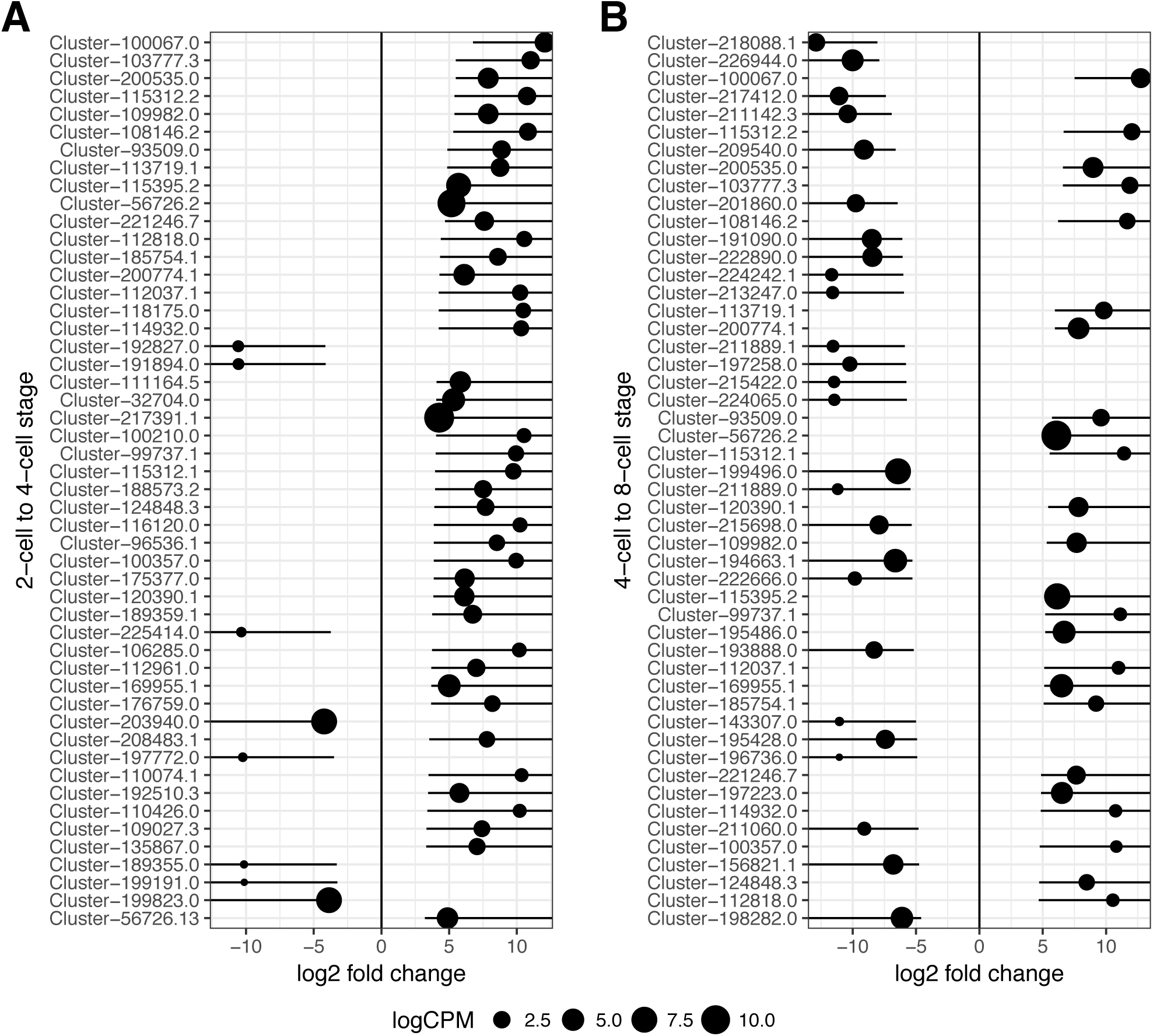
Gene clusters with highest effect sizes (including confidence bounds) for (A) the 2-cell to 4-cell stage and (B) the 4-cell to 8-cell stage; logCPM = log2(counts-per-million).

Differential expression is first detected in the majority of embryonic genes at the 2-cell to 4-cell stage (Figure 7). Effect sizes and confidence intervals were calculated for all DE genes revealing relatively large differences between developmental stages. Genes with the largest effect sizes were predominantly upgregulated at the 2- to 4-cell stage with a more even ratio of upregulated / downregulated genes at the 4- to 8-cell stage (Figure 8). The ratio of total upregulated and downregulated DE genes was similar between developmental stages (Figure 9). This pattern of genome activation in spiny mouse embryos resembles that of the mouse embryo, however the expression of specific genes such as HSP70 (Figure 10F) and the overall profile of transcript expression (Figure 11) share commonalities with the EGA in humans.

**Figure 9:**
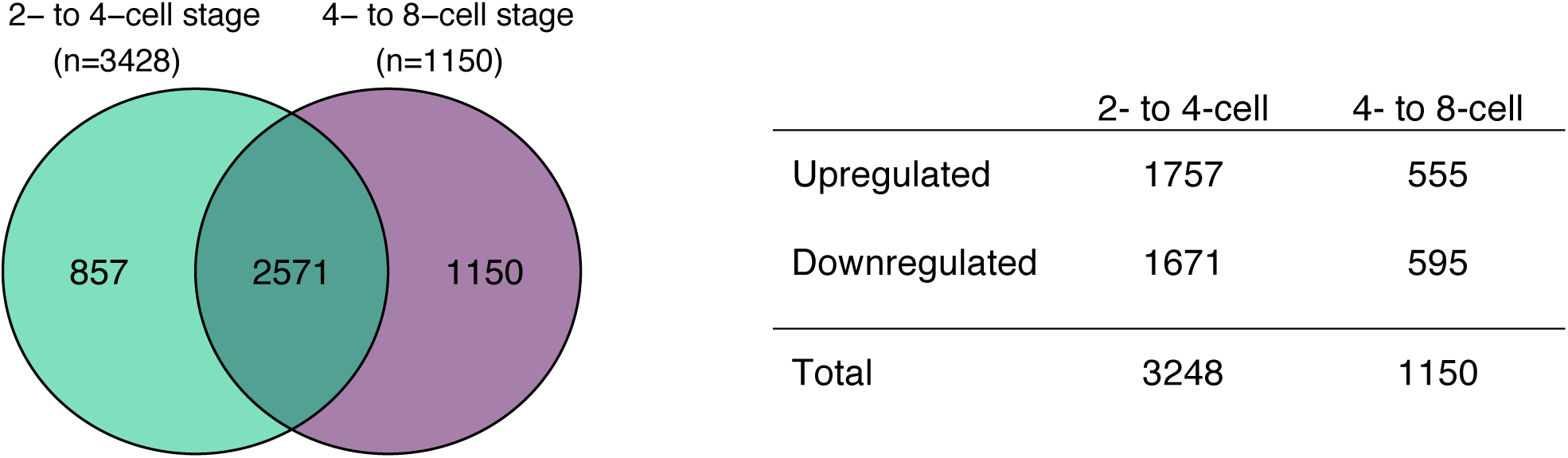
Number of differentially expressed genes at each developmental stage (FDR <0.05). The overlap represents DE genes common to both stages, but first transcribed at the 2- to 4-cell stage. The total number of differentially-expressed genes is further differentiated into upregulated and downregulated genes.

**Figure 10:**
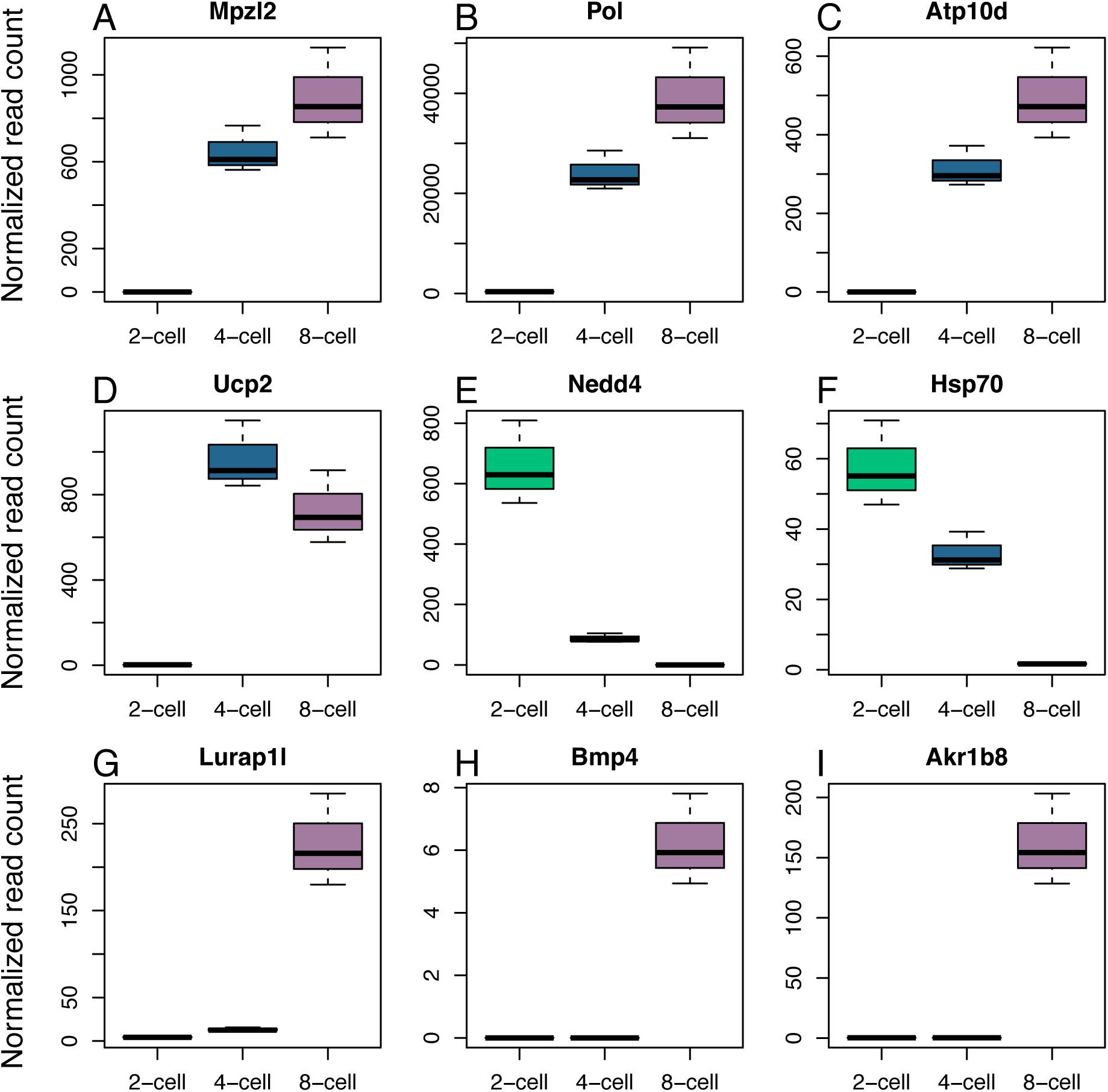
Gene expression profiles for genes-of-interest identified by TopConfects. Expression patterns include increasing expression from the 2-cell to 8-cell stage (A, B & C), high-to-low expression (D, E & F) and expression initiated at the 4-cell to 8-cell stage (G, H & I).

**Figure 11:**
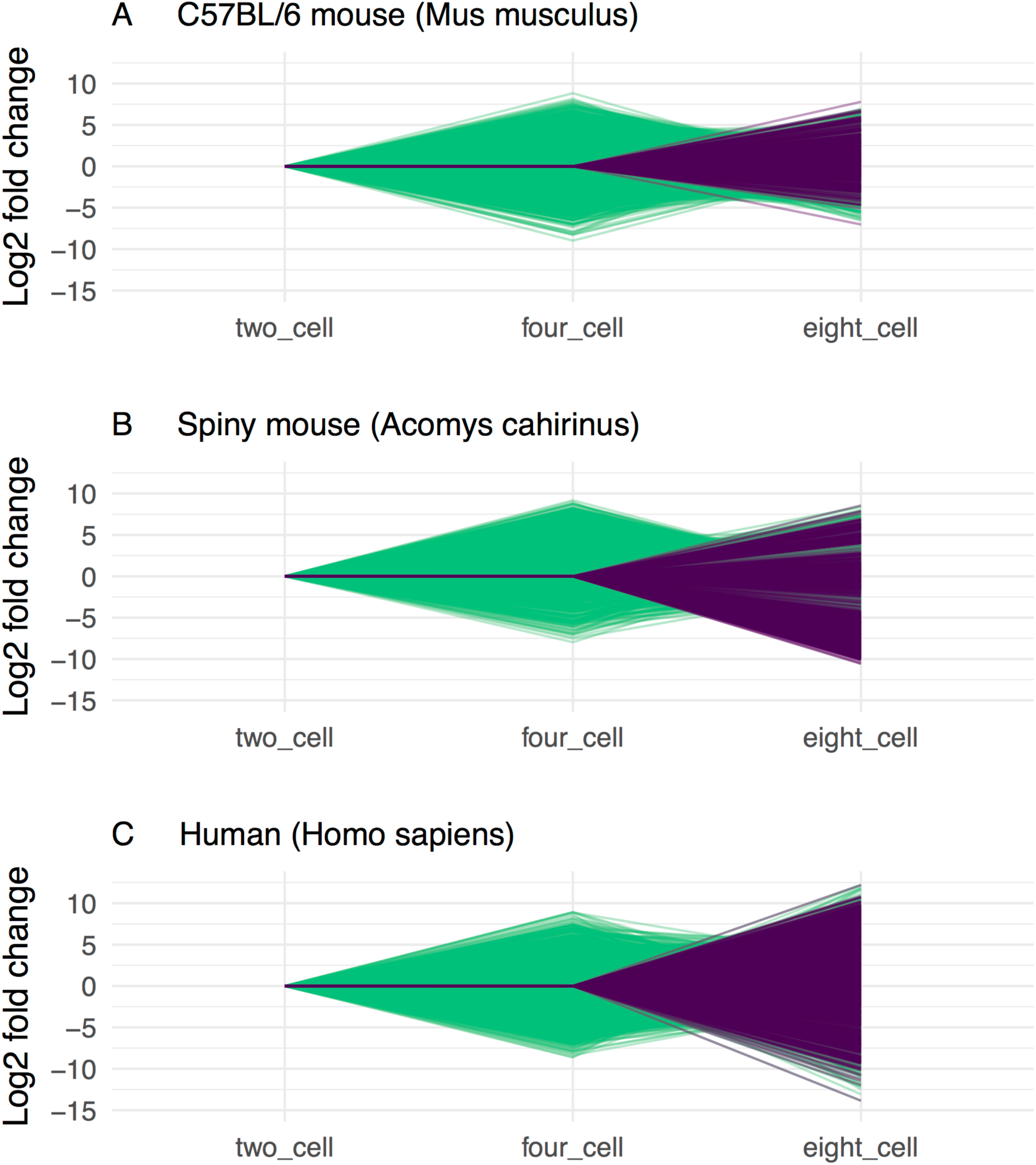
Gene expression profiles for (A) the C57/BL6 ‘common’ mouse, (B) the spiny mouse and (C) for human embryos during the EGA. Genes in which expression is first detected between the 2- to 4-cell stages are represented by green lines. Genes in which expression is first detected between the 4- to 8-cell stage are represented by purple lines. Fewer transcripts are first expressed between the 4- to 8-cell stage in mouse embryos compared to spiny mouse and human embryos. Differences in expression of genes activated at the 4- to 8-cell stage in mice are smaller than spiny mouse and human embryos, displaying less extreme log2 fold changes.

### Profiling gene expression during the EGA

Known differences in specific genes activated during the EGA in mice and humans limit direct comparisons between these species and the spiny mouse, however analysis of overall patterns of transcription are used here to approximate similarities / dissimilarities between species. Gene profiles shown in Figure 10 illustrate different patterns of expression seen in genes of interest identified by the effect size analysis. Except for Hsp70, these genes are not known to play an important role in the EGA in mammals; they are presented to illustrate typical expression patterns seen during the EGA (increasingly high expression, high-to-low expression, and delayed expression until the 4- to 8-cell stage). To determine whether the EGA in the spiny mouse embryo more closely reflects the EGA in human or mouse embryos, profiles were generated for mouse, spiny mouse and human embryos illustrating the pattern of gene expression changes between the 2-, 4- and 8-cell stages for each species (Figure 11). Gene expression changes are less extreme in the mouse embryo, and a smaller number of genes are differentially expressed between the 4- to 8-cell stage compared to spiny mouse and human embryos.

## DISCUSSION

Here we show that the embryonic genome activation (EGA) begins between the 2-cell and 4-cell stages of embryo development in the spiny mouse. This time-point had the greatest number of differentially expressed (DE) transcripts and transcripts for several genes reported to drive the EGA in other mammalian species were identified at this developmental stage for the first time, such as Hsp70 (Bensaude et al., 1983: Figure 10F), Eif4e (Yartseva & Giraldez, 2015), Eif1a (Lindeberg et al., 2004) and Elavl1 (Bell et al., 2008) (Figure S10). The pattern of transcription was similar to other mammals in which the EGA has been characterized (Svoboda, 2017), with massive changes in gene expression occurring within a relatively short time frame. Characteristics used to delineate between the common mouse, spiny mouse and human embryo include the expression of specific genes, the timing of EGA initiation and the ‘burst’ of transcription required for continued development (Richter & Sonenberg, 2005). By these criteria, findings from this study suggest the spiny mouse is a closer model of human embryonic gene expression than the common mouse. This is the first assessment of the spiny mouse for this purpose and these findings warrant further investigation.

An unexpected outcome of this study was sample contamination. Embryo collection was conducted very quickly to minimise the effect of stress on gene transcription and the increased speed of embryo collection resulted in two of the samples becoming compromised. Initial gene expression analysis conducted using the DEGUST web platform (http://degust.erc.monash.edu) revealed this unexpected technical complication. These samples were unable to be salvaged as only ∼30% of the reads they contained aligned to mammalian proteins in the NCBI nr database (Figure S1). This contamination limited our ability to use the *de novo* assembly as a reference for read alignment as a large proportion of the assembled transcripts were found to represent prokaryotic sequences rather than spiny mouse sequences. Use of this assembly would have resulted in erroneous quantification of gene expression. Extracting read-supported embryo-specific transcripts from the ‘embryo’ assembly and adding them to the reference spiny mouse transcriptome (Mamrot et al., 2017) was a successful solution for avoiding transcripts derived from prokaryotic organisms. Several genes that are only expressed during early embryo development can now be found in the reference transcriptome, such as Oct3/4, Nanog, Oobox and H1foo. The updated assembly has been uploaded to a permanent data repository (https://doi.org/10.5281/zenodo.1188364) and is accessible via our BLAST search website: http://spinymouse.erc.monash.edu/sequenceserver/ (“Trinity_v2.3.2_plus_embryo-specific_transcripts”) (Priyam et al., 2015). This approach significantly improved gene-level resolution and improved the reference transcriptome for future applications.

A downstream effect of sample contamination was reduced statistical power. Our preliminary power calculation predicted 4 samples per group would be required to accurately quantify differences in gene expression between developmental stages (Figure S2). Exclusion of two samples reduced our ability to resolve DE genes, however the parameters used for the initial power calculation were found to be relatively conservative and the analysis was modified to mitigate against this confounding factor. Quasi-likelihood F-tests were used to establish differential expression (rather than likelihood ratio tests) to gain stricter error rate control by accounting for uncertainty in the original dispersion estimate (Chen et al., 2016). In addition, transcript expression was analysed at the ‘gene’ level to avoid potential biases previously reported in transcript-level analyses (Kanitz et al., 2015; Leshkowitz et al., 2016; Williams et al., 2017). This modified workflow was effective in identifying a large number of DE genes, however the total number of DE genes reported for each timepoint are likely to be underestimated. Using an unadjusted p-value (p=0.05) as a cutoff for statistical significance, rather than adjusting the p-value to reduce the false discovery rate (FDR=0.05), provides an indication of genes that may have been detected as differentially expressed given full experimental power (Figures 7B & 7D). This suggests the number of genes upregulated between the 2-cell and 4-cell stage and number of genes downregulated between the 4-cell and 8-cell stage are likely greater than reported.

The presence of rRNA reads was another unexpected outcome. Rather than sequence poly(A)+ RNA, we depleted rRNA using custom designed depletion probes manufactured by NuGEN (formerly known as Insert Dependent Adaptor Cleavage “InDA-C” probes) to obtain non-coding RNA transcripts and partially-degraded maternally-inherited transcripts in addition to mRNA (Bush et al., 2017; Schuierer et al., 2017). This approach was partially successful. Greater than 80% of total RNA is composed of rRNA in preimplantation embryos (Bush et al., 2017; O’Neil et al., 2013; Piko & Clegg, 1982), so levels detected in our samples (∼30-40%: Figure 1) suggest the AnyDeplete rRNA probes worked, but were not fully effective. There are several potential explanations for this result; the most likely explanation is that our AnyDeplete probes were designed and tested using spiny mouse RNA-Seq derived transcripts whereas AnyDeplete probes are typically designed using a reference genome (a spiny mouse genome is not yet publicly available). The impact of rRNA levels on the ability to detect relative abundance of protein-coding RNA transcripts in preimplantation embryos is unknown.

Protein-coding genes known to regulate early development in mammalian embryos were detected at the 2- to 4-cell stage in the spiny mouse. These genes, including Yap1, RNA polymerase II, E3 ubiquitin-protein ligase, and the eukaryotic initiation factor family of transcripts have been implicated in the EGA in humans through various mechanisms of action (Ge, 2017; Svoboda 2017). One of the first proteins transcribed in mammalian embryos is the Heat Shock Protein 70kDa (known as Hsp70 / Hspa1a) (Bensaude et al., 1983). This protein performs several roles during the MZT, such as establishing chromatin structure, genome stability, and chaperoning O-linked glycosylated proteins into the cell nucleus (Abane & Mezger, 2010; Guinez et al., 2005; Nagaraj et al., 2017). In spiny mouse embryos Hsp70 expression is relatively high at the 2-cell stage followed by decreasing expression at the 4- and 8-cell stages. High expression of this gene during the first ‘wave’ of the EGA has been shown in many species, including the mouse, bovine and human embryo (Bettegowda et al., 2007; Christians et al., 1997; Lelièvre et al., 2017). Early transcription of HSP70 is crucial for successful cell cleavage and continued development, with compromised gene expression and protein levels correlated with embryo cell blocks. This pathway provides a potential target for understanding and overcoming the 4-cell block in the spiny mouse.

Direct comparison of DE gene sets between mouse, spiny mouse and human embryos at these stages of development was not conducted due to poor inter-species consensus reported by others (e.g. Heyn et al., 2014; Xie et al., 2010), however specific genes directly implicated in the EGA were investigated (Figure S10). Overall variation in EGA-related gene expression was found between the mouse, spiny mouse and human, with the results for these gene sof interest (Eif4e, Elavl1, Pou5f1, Eif1a) representing typical inter-species differences. Although differences were identified in this study between the mouse, spiny mouse and human, further efforts to replicate and reproduce these results would increase the likelihood that these findings represent differences in the underlying mechanisms driving the EGA, rather than other factors. A more robust inter-species comparison of the EGA is the overall changes in gene expression patterns during these early developmental stages (Figure 11). This comparison revealed a closer relationship between spiny mouse and human embryos, compared to the common mouse, with a greater number of genes first expressed at the 4- to 8-cell stage and a larger range of expression changes during this period of development. These findings support use of the spiny mouse (*Acomys cahirinus*) as a model of the human EGA.

In conclusion, anatomy and physiology varies between all animal models of human reproduction and development. Primates are arguably the most accurate representation of human physiology, with similar anatomy and endocrine profiles, however ethical and logistical constraints limit their usefulness for basic research. Rodents offer an attractive alternative, as they have short breeding intervals and their anatomy and physiology has been comprehensively studied. Despite the advantages, translation of findings from mice to humans is not always successful, suggesting the common mouse may not be the best model for early human development. Conspicuously, the absence of a menstrual cycle in the common mouse is associated with key differences in how embryos are formed and develop. Here, we aimed to investigate the spiny mouse and assess its usefulness for modelling early human embryonic gene transcription. Methodological limitations impacted our ability to comprehensively address this aim, however the novel findings reported here support further investigation into other aspects of embryology in this species. Future directions for this work include further sequencing of spiny mouse embryos at the zygote, 16-cell stage, morula and blastocyst stages, and use of this RNA-Seq dataset to investigate the conditions required to overcome the 4-cell block in the spiny mouse embryo to facilitate further comparison of embryos developed *in vitro* and *in vivo* in this species.

## Acknowledgements

We thank Vivien Vasic, Trevor Wilson and members of the MHTP Genomics platform for conducting the challenging library prep and RNA-Seq. Tony Papenfuss for providing access to WEHI facilities. David Powell for his assistance with the spiny mouse blast database website and for use of Monash Bioinformatics Platform resources. David Walker for his ongoing support. Ashleigh Clark and Nadia Bellofiore for maintaining the spiny mouse colony.

## Author contributions

JM and HD designed the study with advice from DKG and PTS. JM collected and prepared samples for sequencing, assembled and analysed the sequencing data, prepared figures and wrote the manuscript. All authors read and approved the final version of the manuscript.

## Supplementary figures and tables

**Figure S1:**
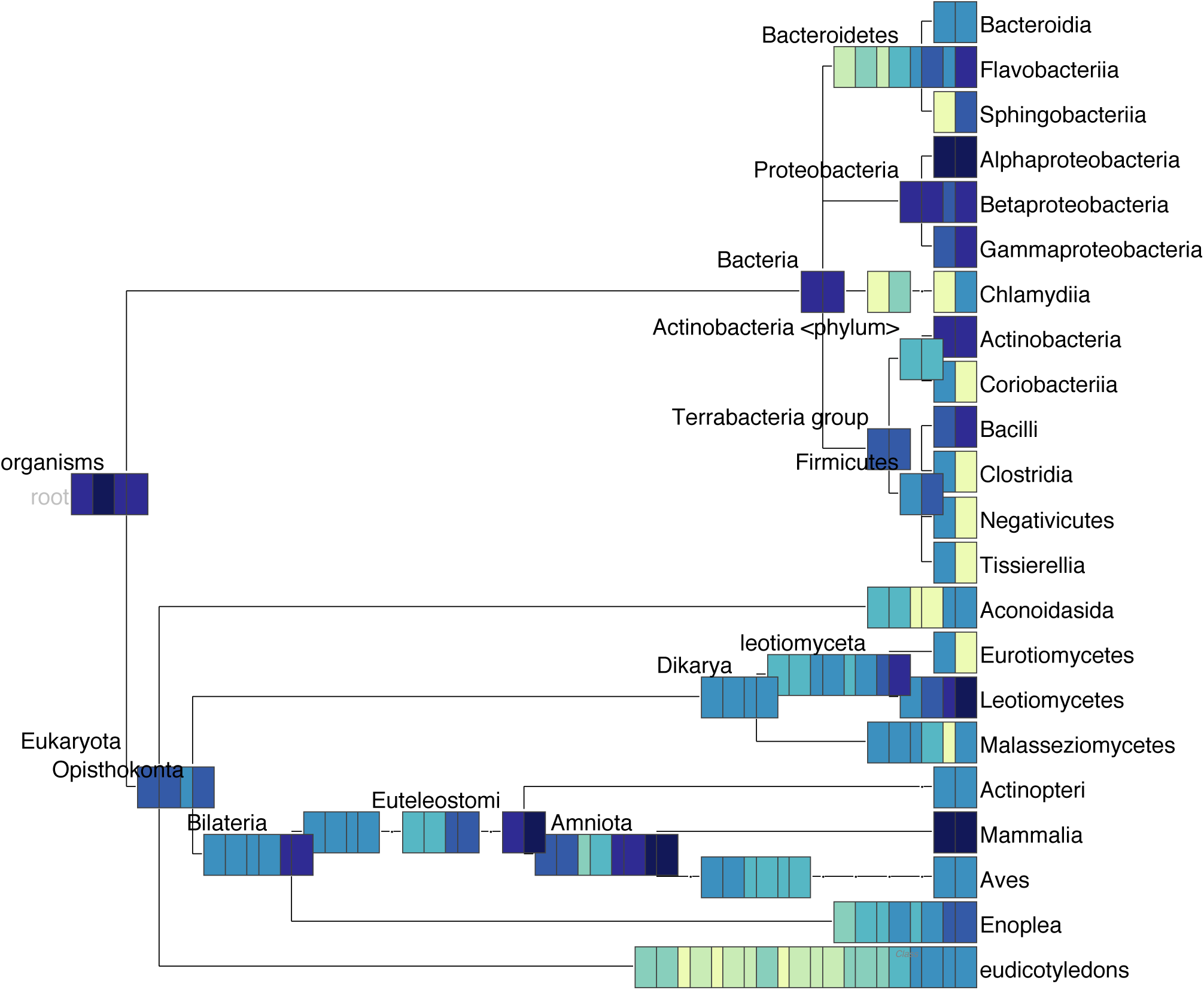
Metatranscriptomic analysis of Trinity-normalized reads in samples 2-cell_C and 8-cell_A illustrating read alignment to eukaryotic and prokaryotic taxa within the NCBI nr protein database (“non-redundant” proteins; n=4,348,972). Read alignments were summarised at the Class level using MEGAN6 implementing the ‘Blues’ colour scale: a higher proportion of aligned reads is represented by a darker colour (highest number of reads aligned per taxonomic group = “Midnight Blue”). Within these two contaminated samples ∼30% of total reads aligned to Mammalia, ∼30% aligned to Alphaproteobacteria, and ∼40% were spread across other Classes as indicated.

**Figure S2:**
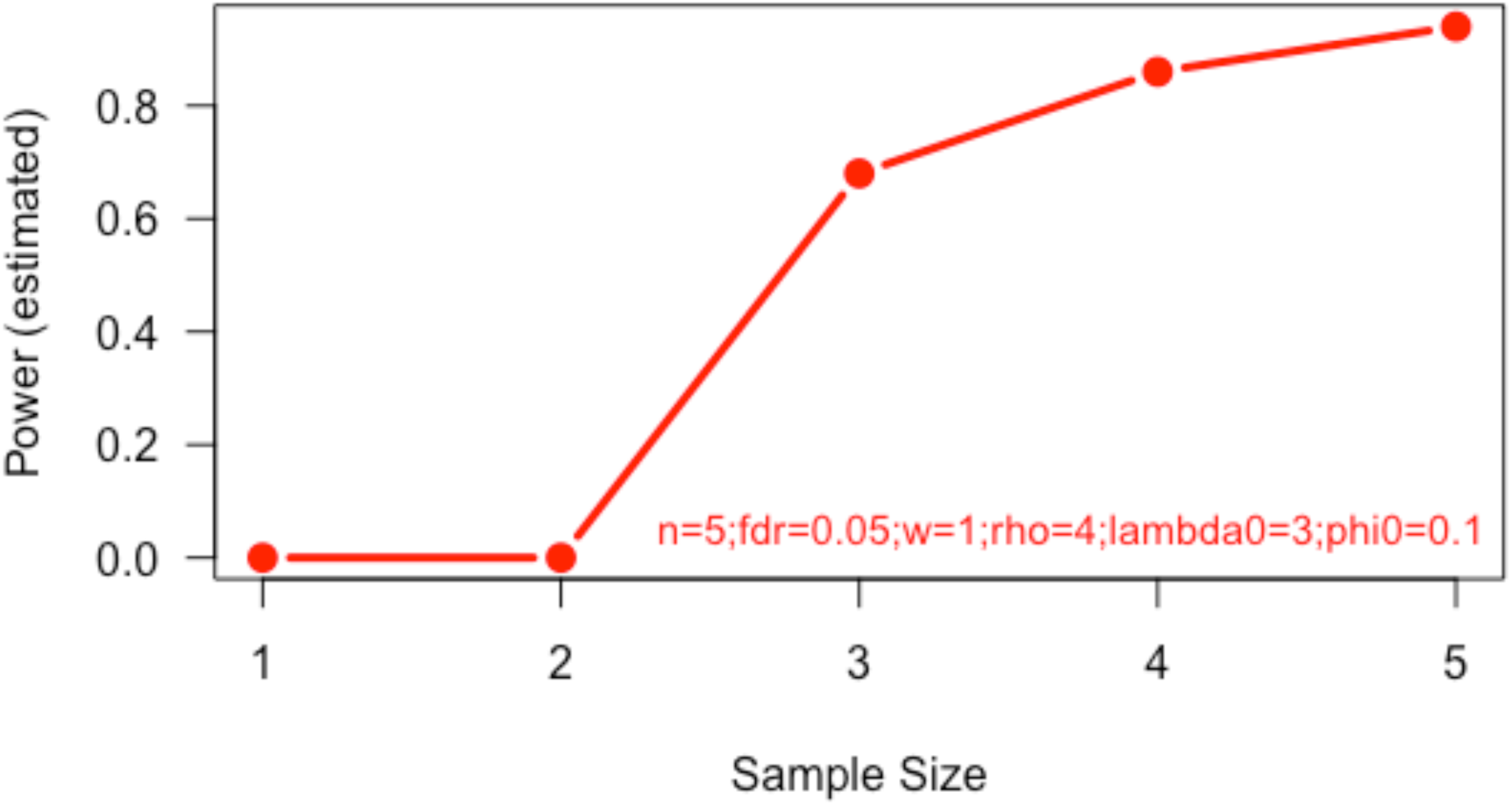
Power estimates for various sample sizes. Parameters represent expected values based on past / similar experiments. “w”: expected normalization factor for sample groups (a value of 1 representing approximately equal read counts across sample groups). “rho”: fold change required for significance, “FC=4” => log_2_(FC)=2. “lambda0”: anticipated average read count per sample (actual values were higher than predicted: Figure 2); “phi0”: average dispersion across samples (actual dispersion value was slightly lower than expected). With the parameters specified, n=4 in each group is recommended to achieve power >0.8.

**Figure S3:**
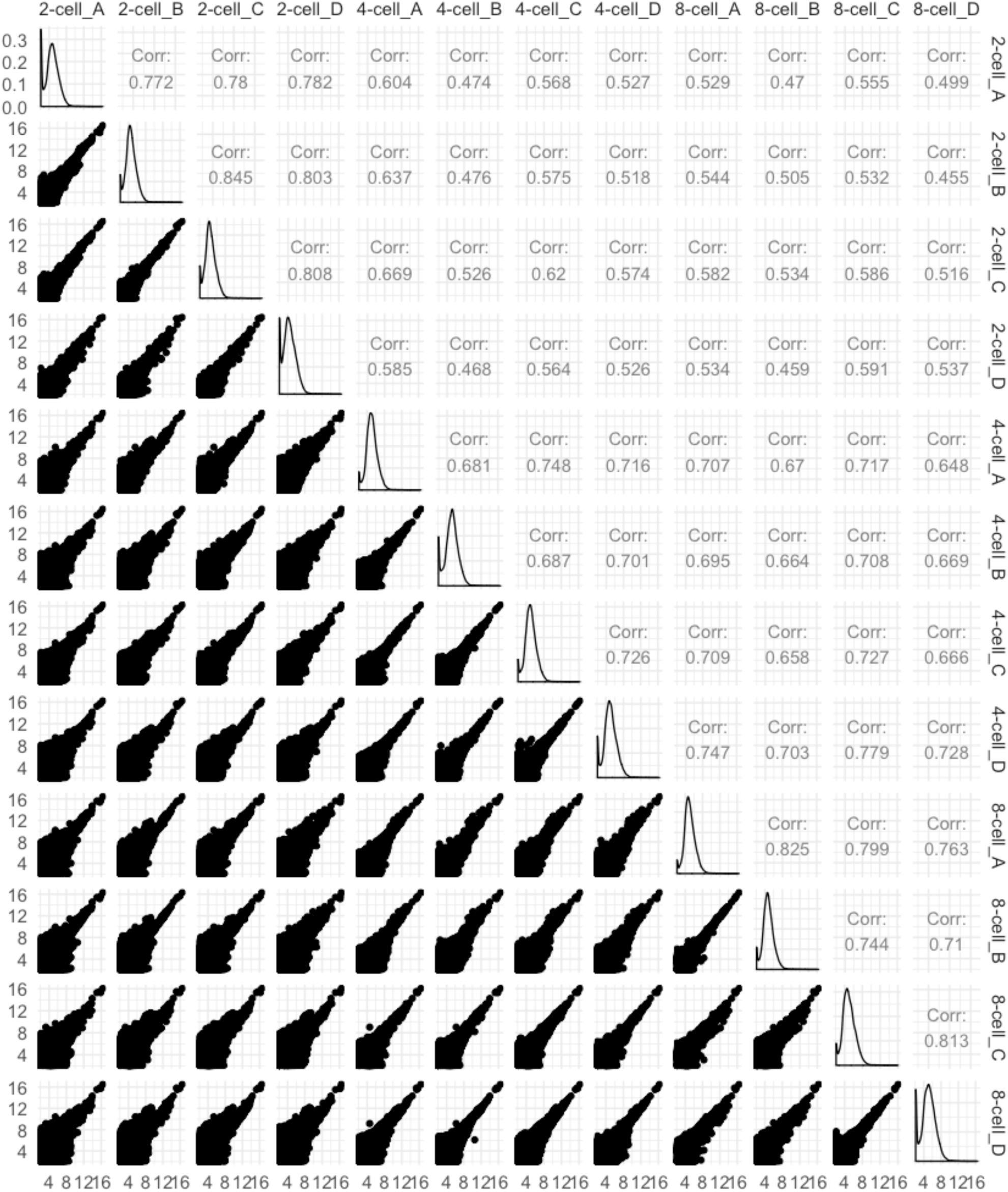
Correlation matrix for all samples (n=12). Spearman correlation values (upper right), distribution (diagonal) and concordance (lower left) of gene cluster abundance are illustrated.

**Figure S4:**
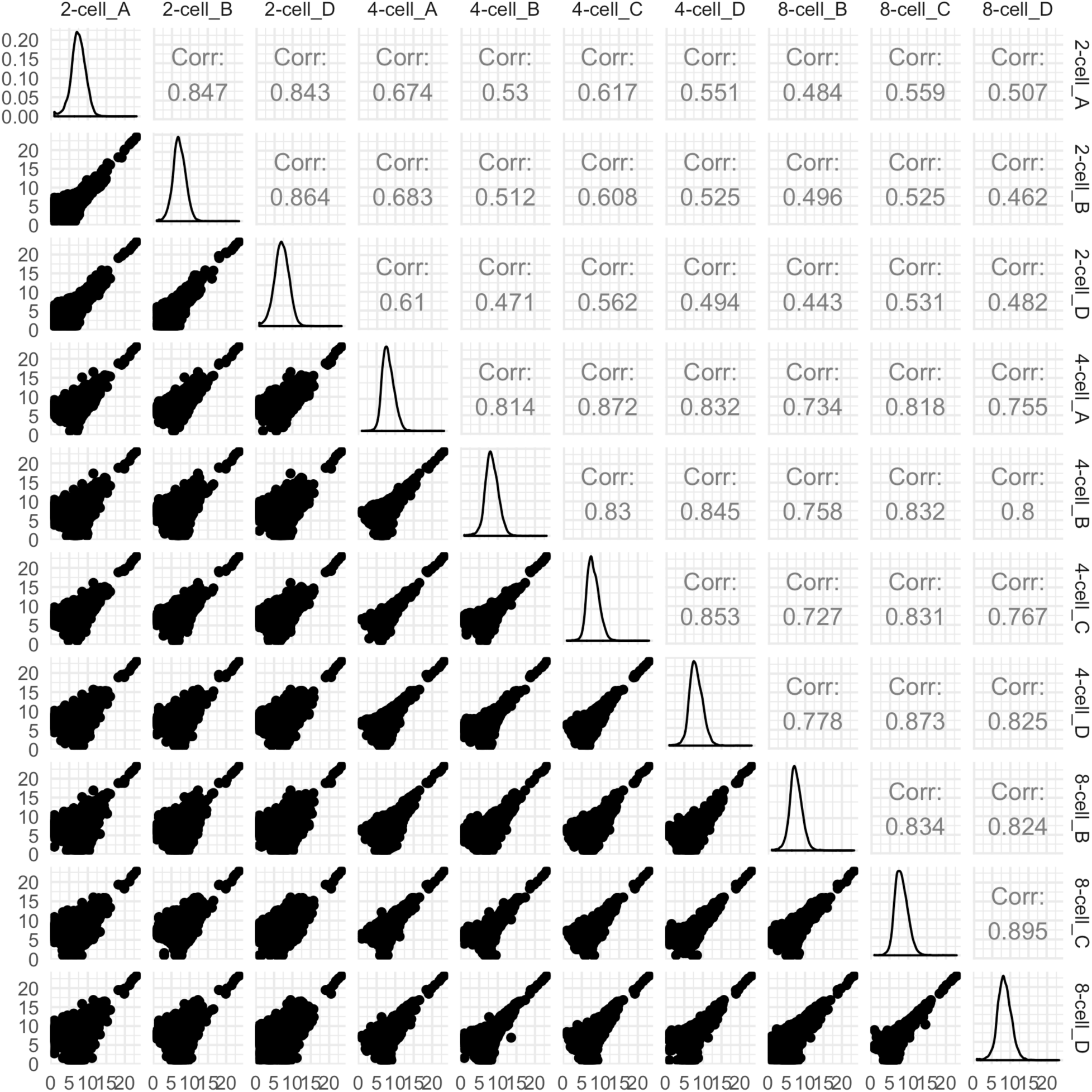
Correlation matrix for uncontaminated samples (n=10). Spearman correlation values (upper right), distribution (diagonal) and concordance (lower left) of gene cluster abundance are illustrated.

**Figure S5:**
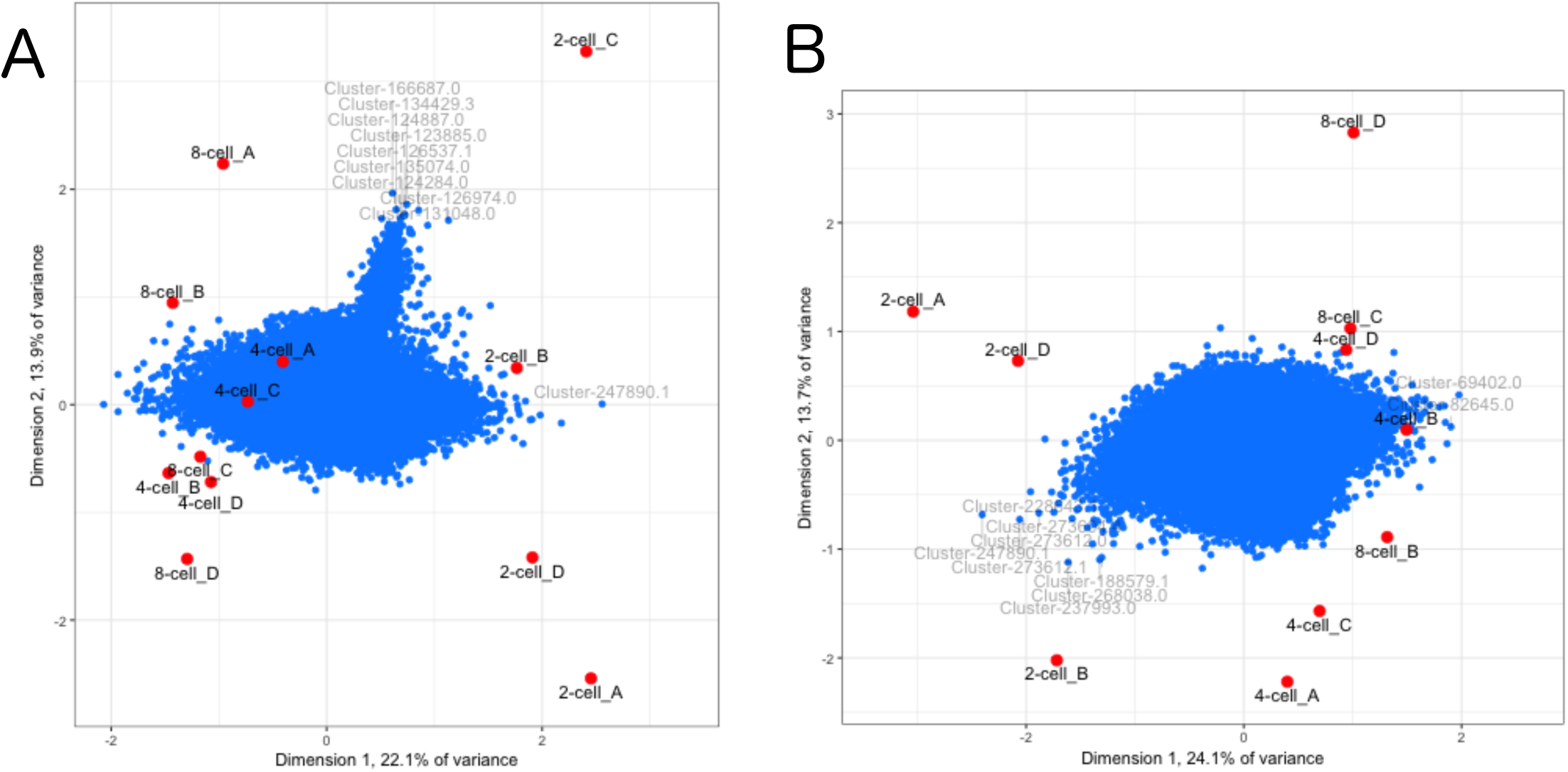
Singular Value Decomposition (SVD) biplots of gene expression per sample. Blue dots represent gene expression values and red dots represent samples, with (A) contaminated samples included, and (B) with contaminated samples excluded. The top 10 differentially expressed genes are labelled in each plot. Many of the top DE genes in (A) correspond to prokaryotic taxa (9/10). In comparison, after the contaminated samples were excluded in (B) the top DE genes all correspond to mammalian taxa.

**Figure S6:**
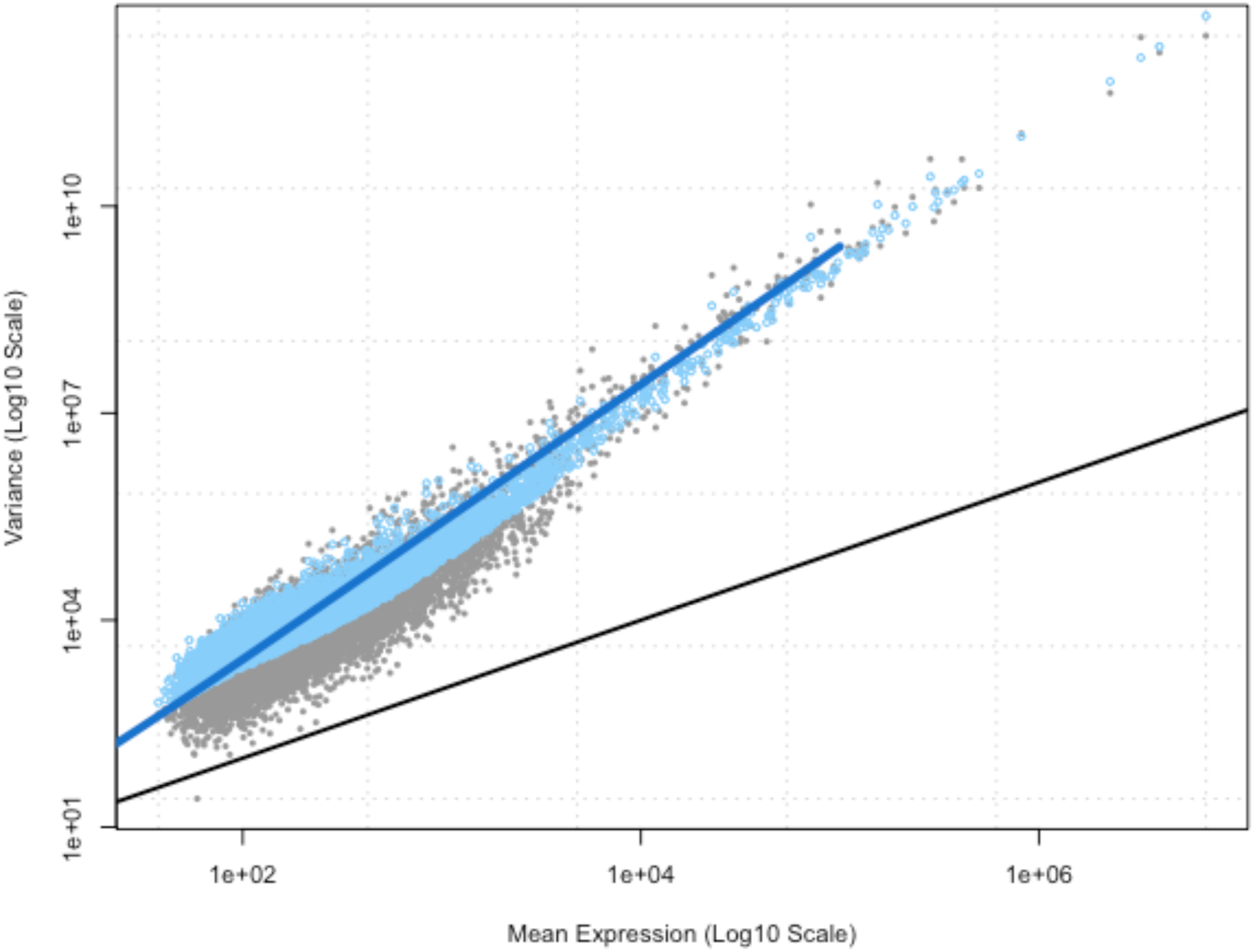
Fit of the edgeR negative binomial distribution to gene counts.

**Figure S7:**
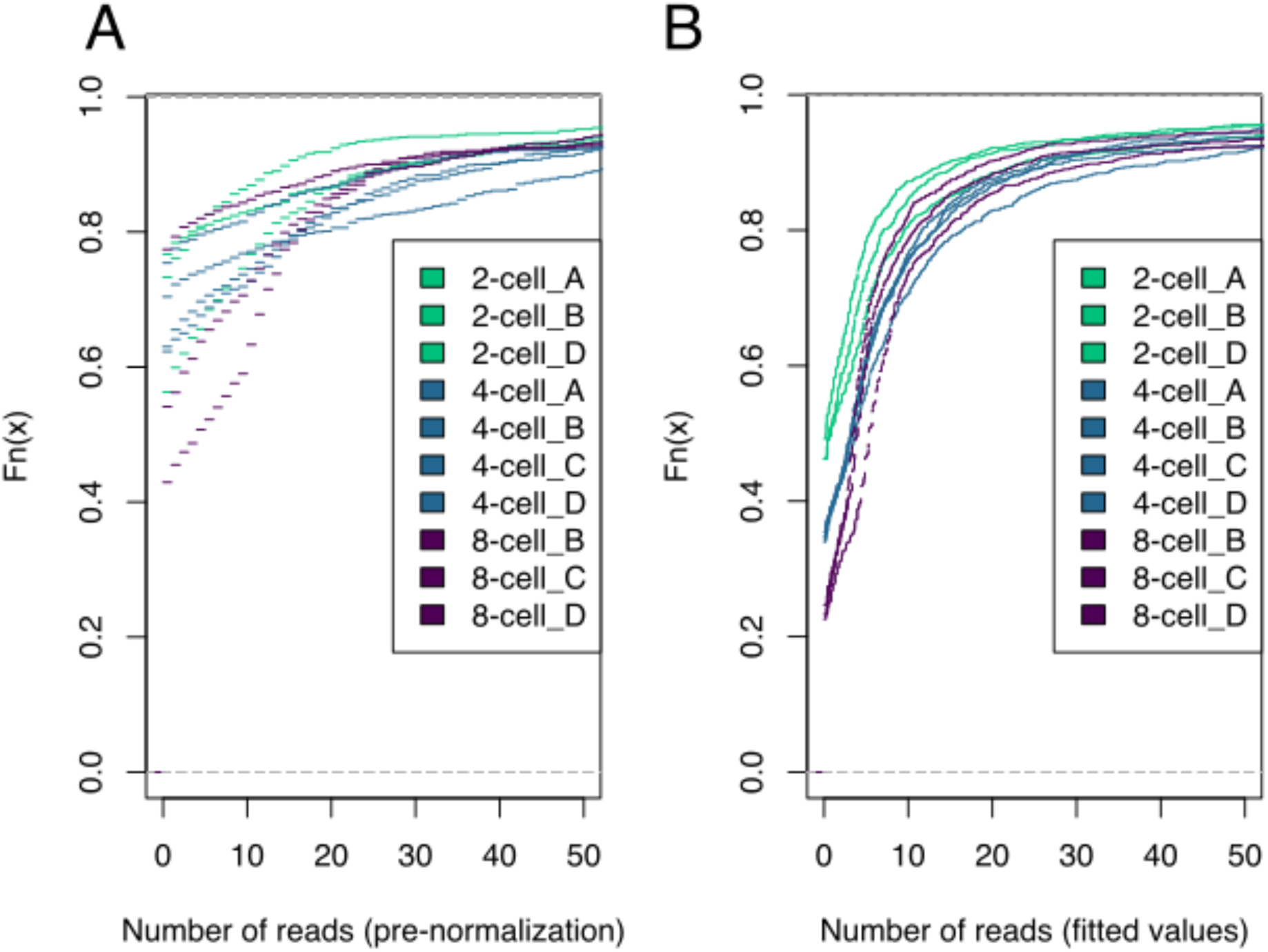
Multiple empirical cumulative distribution of reads for each sample. (A) Read counts for all genes prior to normalization. (B) Normalization and fitting using the negative binomial model improved grouping by developmental stage, especially for below-average read counts.

**Figure S8:**
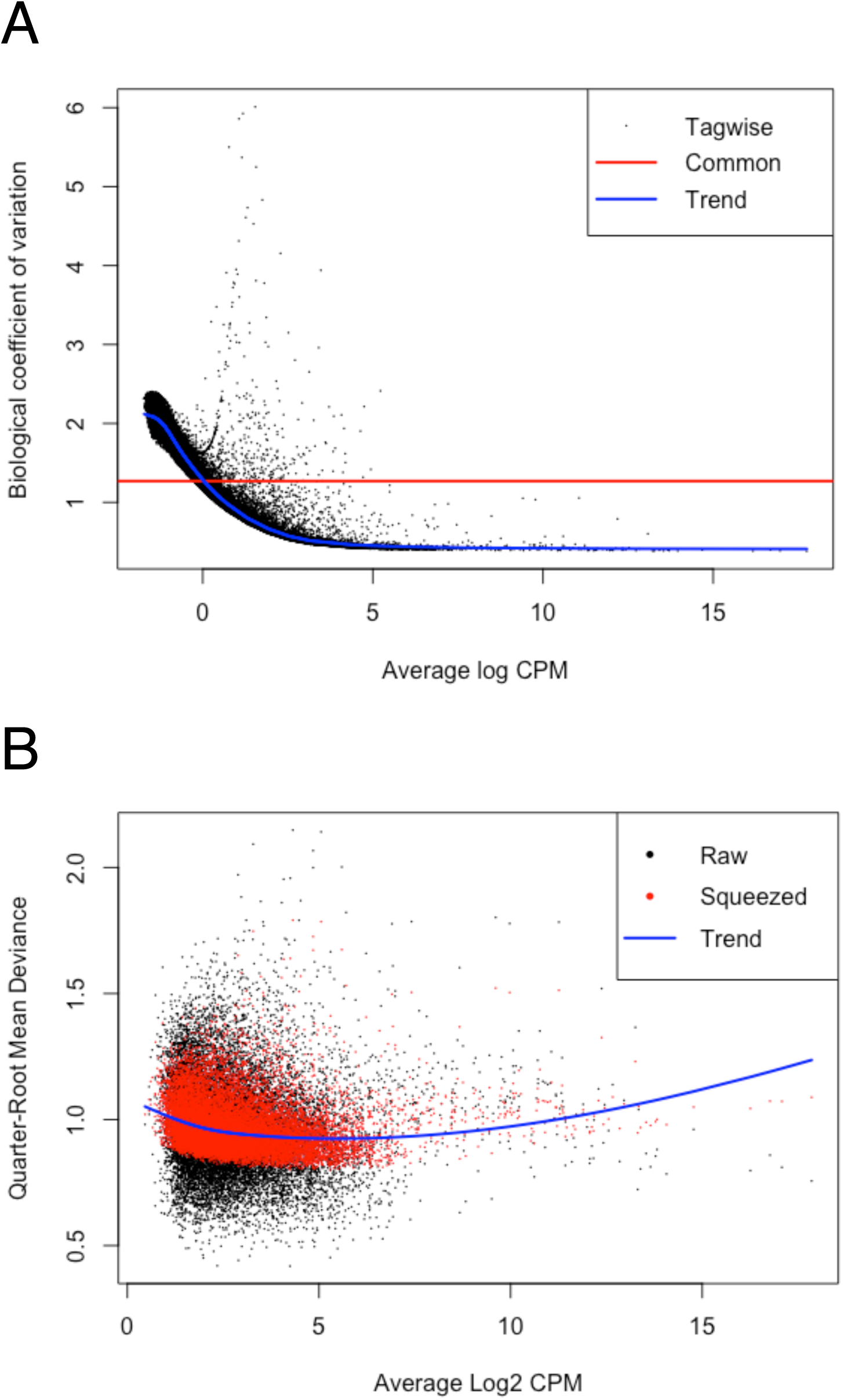
Scatterplots illustrating (A) the biological coefficient of variation and (B) the quarter-root of the quasi-likelihood dispersions for all genes. cpm=counts-per-million.

**Figure S9:**
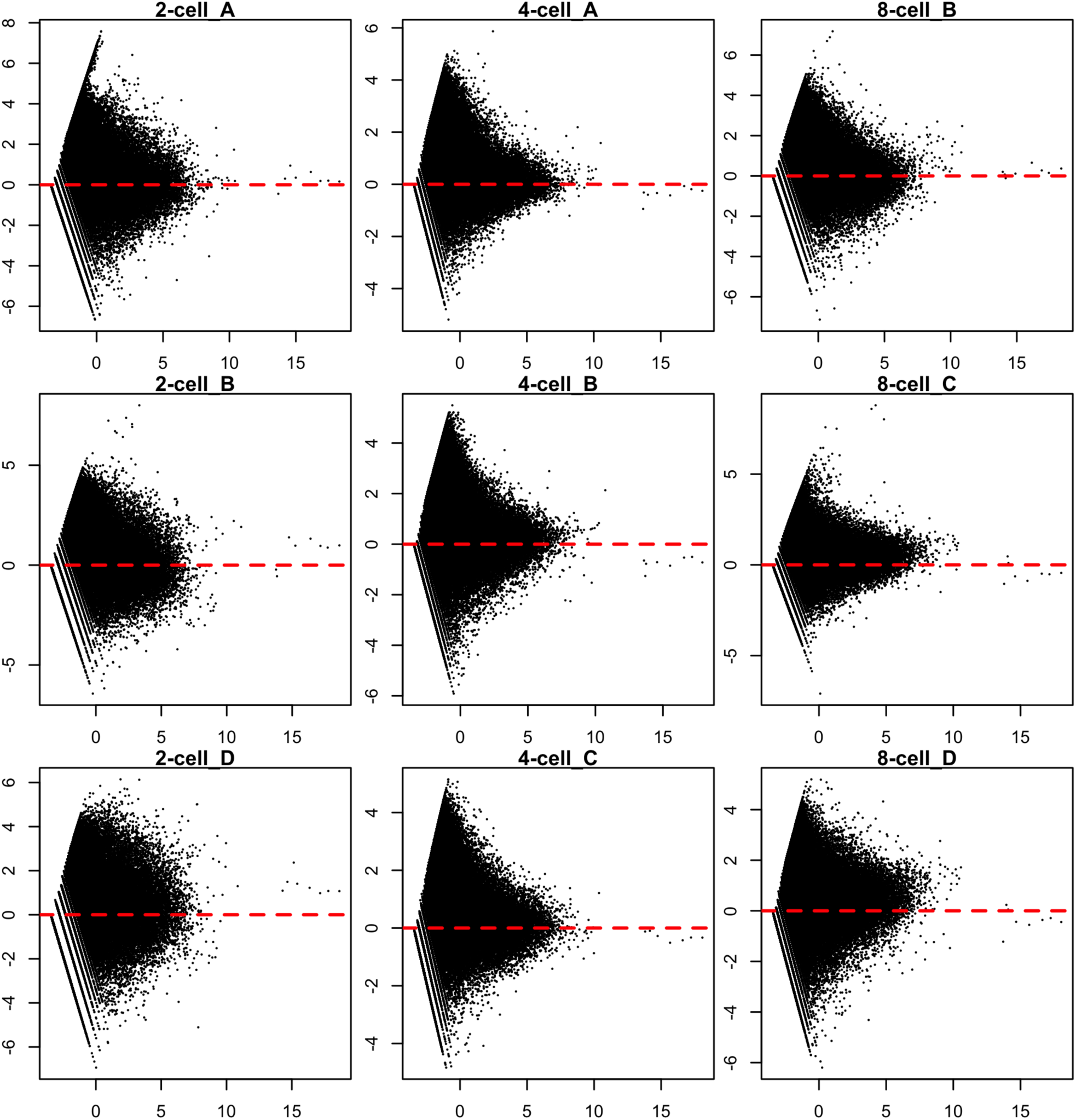
Mean-Difference (MD) plots comparing each uncontaminated sample to an artificial reference library constructed from the average of all other samples. Sample 4-cell_D was included in the analysis but excluded from this figure for readability; full figure with all samples: https://doi.org/10.4225/03/5a9531283d103. Positive skew in samples (eg 8-cell_C and 8-cell_D) corresponds to greater variation in TMM normalization factors (Table S2).

**Figure S10:**
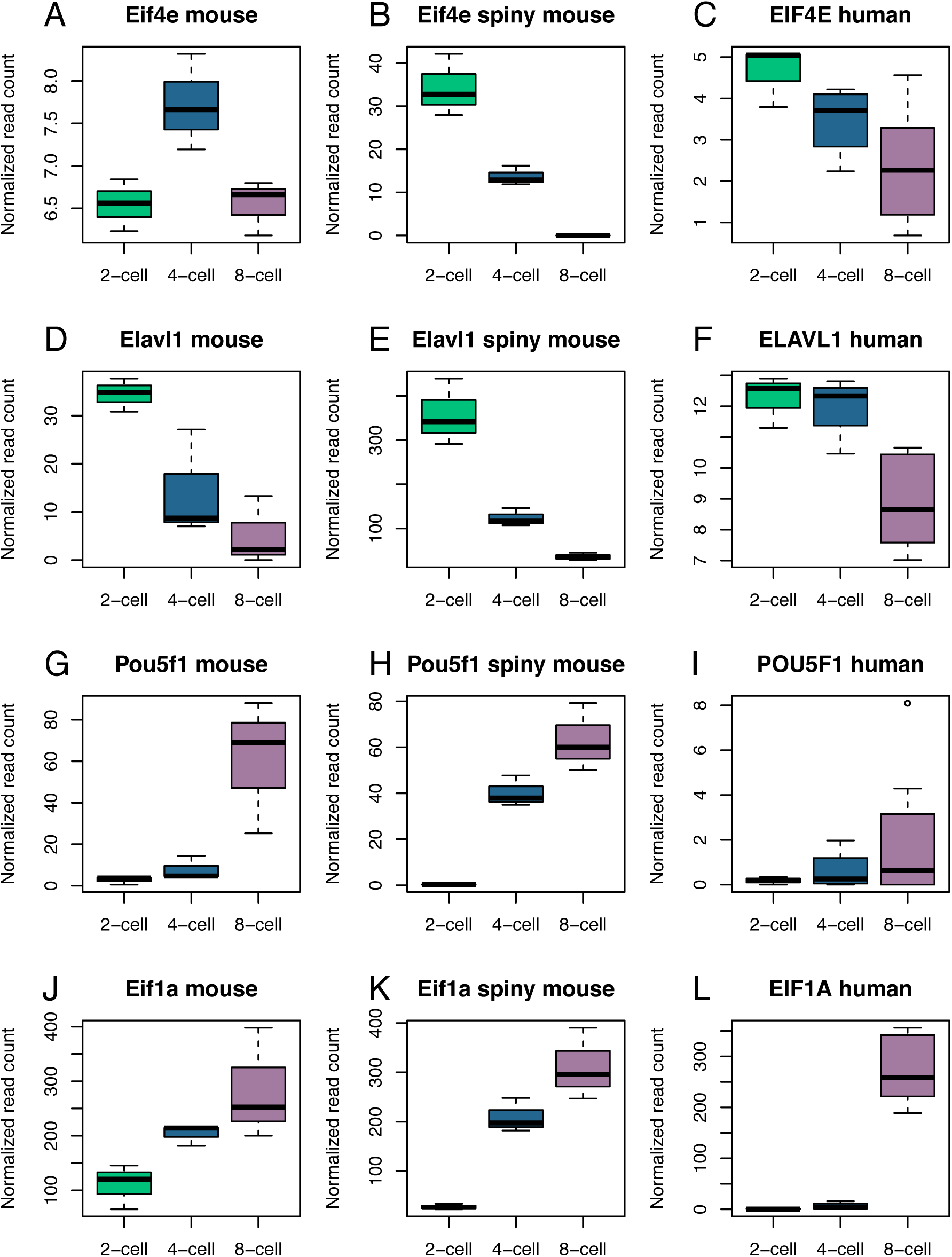
Expression of select genes in common mouse, spiny mouse and human embryos during the 2-cell, 4-cell and 8-cell stages of development. Eukaryotic translation initiation factor 4E (EIF4E) is a key component of the translation machinery and a known driver of genome activation in mammals. ELAV like RNA binding protein 1 (ELAVL1) is an RNA stabilizer involved in maternally-inherited transcript clearance. POU Class 5 Homeobox 1 (POU5F1), also known as OCT3/4, is a key regulator of pluripotency with highest expression at the morula and blastocyst stages. Eukaryotic translation initiation factor 1A (EIF1A) is required for protein biosynthesis and an increase in expression occurs during the EGA.

**Supplementary Table 1:**
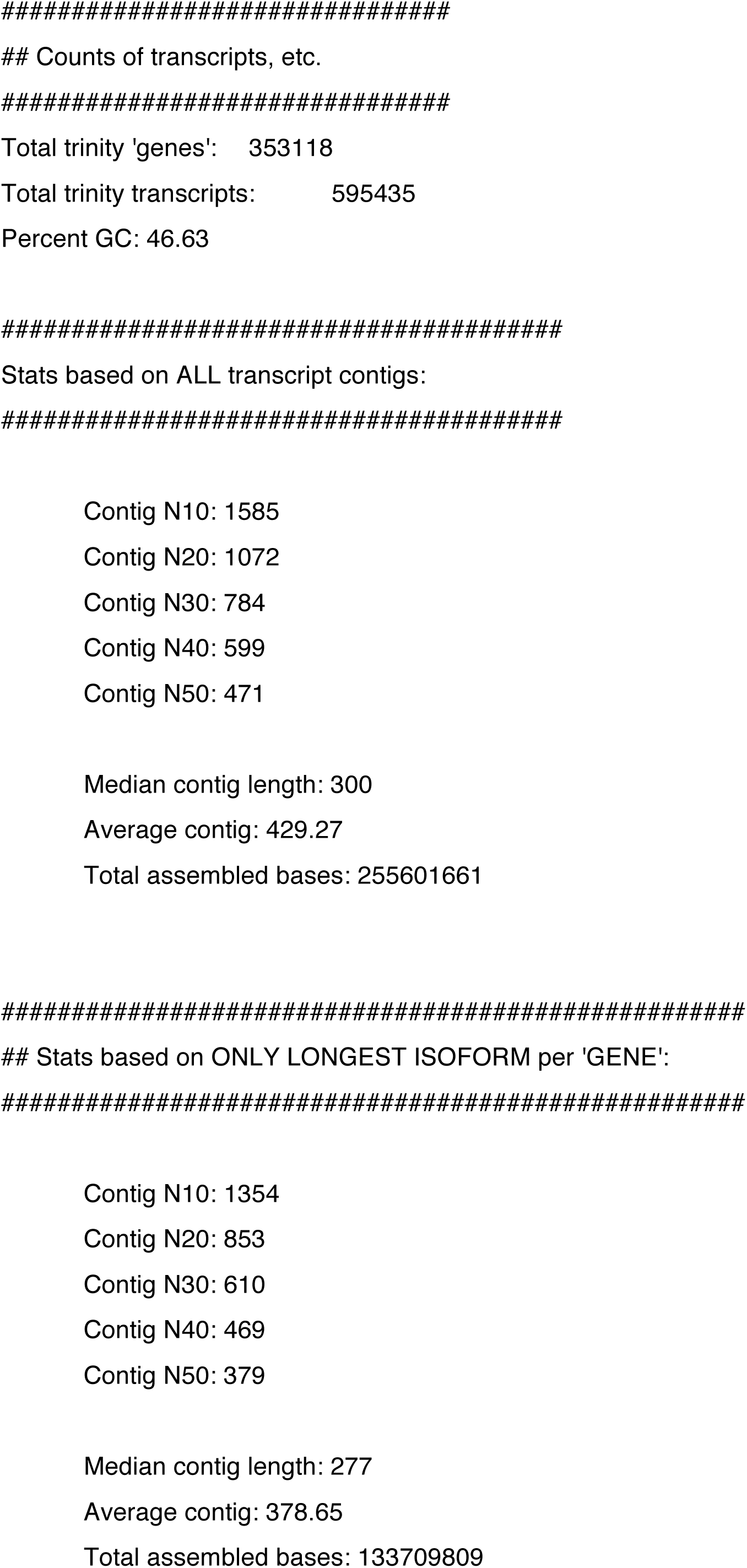
Statistics for Trinity transcriptome assembly (output from TrinityStats.pl).

**Supplementary Table 2:** TMM-normalized library sizes

| Sample | Library size | Adjusted library size |
| --- | --- | --- |
| 2cell_A | 30584677 | 30523713.55 |
| 2cell_B | 32974138 | 24386500.70 |
| 2cell_C | 37049595 | 22227903.12 |
| 2cell_D | 35508389 | 20709266.29 |
| 4cell_A | 31371210 | 38551172.77 |
| 4cell_B | 32900876 | 48943239.95 |
| 4cell_C | 28156913 | 36219884.07 |
| 4cell_D | 29500819 | 39510735.95 |
| 8cell_A | 34693598 | 31935445.77 |
| 8cell_B | 29556455 | 28958706.22 |
| 8cell_C | 26386705 | 29998846.68 |
| 8cell_D | 34873989 | 41815650.37 |

